# Optimization of the glycerol production from *Dunaliella tertiolecta* and *Dunaliella* isolates

**DOI:** 10.1101/2023.09.18.558238

**Authors:** Linda Keil, Jonas Breitsameter, Bernhard Rieger, Daniel Garbe, Thomas Brück

## Abstract

The salt tolerant, marine microalgae *Dunaliella tertiolecta* is reported to generate significant amounts of intracellular glycerol as an osmoprotectant under high salt conditions. Several studies have identified light color and -intensity, nitrogen- and phosphate limitation, as well as salt concentration as the most inducing factors impacting glycerol productivity. This study aims to optimize glycerol production by investigating these reported factors singularly and in combination to improve the glycerol product titer. The reference strain *D. tertiolecta* as well as new *Dunaliella* isolates were evaluated in this study. The results demonstrate that cultivation with white light of an intensity between 500 and 2000 μmol m^−2^ s^−1^ as opposed to 100 μmol m^−2^ s^−1^ achieves higher biomass and thereby a higher product titer. Moreover, cultivation in 1.5 M NaCl followed by an increase to 3 M NaCl resulted in hyperosmotic stress conditions, providing the highest glycerol titer. Under these optimal light intensity and salt conditions, the glycerol titer of *D. tertiolecta* could be doubled to 0.79 mg mL^-1^. Furthermore, under the same conditions glycerol extracts from new *Dunaliella* isolates did provide up to 0.94 mg mL^-1^. Highly pure glycerol obtained under optimal production conditions has widespread applications e.g., in the pharmaceutical industry or the production of sustainable carbon fibers. Moreover, it can be labeled as halal.

## Introduction

Glycerol is an essential raw material to produce various valuable products such as pharmaceuticals, food, cosmetics, paints and textiles [1–4]. The glycerol market size reached over USD 2 billion in 2021 and is expected to increase annually by over 7% from 2022 to 2030 [5]. Glycerol will be an important compound for high value chemicals and biopolymers in the future [6]. As a promising precursor for ‘green chemicals’, glycerol has the chance to replace several fossil-based chemicals such as ethylene and propylene glycols, 2,3-butanediol, and acrolein [6–10]. Additionally, glycerol constitutes as a precursor for polyacrylonitrile (PAN), which is one of the most important polymers used in industry [11]. PAN in turn is the precursor for many products, such as resins, plastics and acrylic fibers [12, 13], as well as for carbon fibers [14, 15]. (Poly-) Acrylonitrile is synthetized *via* the SOHIO process [16], which is a fossil-based process *via* ammoxidation of propene [17] and comes along with high carbon dioxide emissions. A less CO_2_ emitting process is the conversion of glycerol to acrylonitrile, especially if bio-based glycerol serves as raw material [13].

Large demands of environmental friendly glycerol are covered through bio-fuel production which generates glycerol as a by-product [13, 18]. Beyond the high prices and availability fluctuations [19], there are many impurities in biofuel [19], which raise the purification costs as complex purification processes are needed [20, 21]. The application in the field of pharmacology and food requires a glycerol purity greater than 99.7%. However, commonly used purification methods of the biofuel-derived glycerol do not achieve the needed purity [22]. Additionally, bio-fuel production utilizes enormous agricultural areas necessary to nourish the world population [23, 24]. Therefore, a sustainable and less area-intensive production process for the extensively required raw material glycerol is of great importance [18, 25]. Used cooking oil might be a sustainable alternative for the bio-fuel and glycerol production [26], but also does not reach a glycerol purify higher than 99.7% [22]. Furthermore, the resulting glycerol is not necessarily halal, which is needed for pharmaceutical and food application with halal quality [27].

Microalgae are an appropriate alternative that allow the environmental friendly synthesis of glycerol [23], as they are unicellular aquatic microorganisms, that perform photosynthesis and thereby convert sunlight, the greenhouse gas CO_2_, and inorganic nutrient salts into biomass and glycerol [28–31]. Compared to terrestrial plants, microalgae grow up to ten times faster [32, 33] and photosynthesize about ten times as efficiently as plants do, owing to their energy efficient photosynthetic apparatus [34, 35]. Moreover, it is feasible to cultivate microalgae on wasteland using saline or waste water [36, 37] with the result that they do not compete with agricultural food production. These advantages make microalgae a suitable biotechnological tool to efficiently fix CO_2_ and convert it into sustainable high-value products [38, 39].

*Dunaliella* species, for example, is a strain that is already cultivated in large industry scale to produce β-carotene [40]. Generally, it is very salt-tolerant and is able to grow with salt concentrations up to saturation [41]. The cells lack a rigid polysaccharide cell wall and are only bounded by a cytoplasmic membrane [42]. This allows the algae to quickly adapt to hyperosmotic changes by reducing the size of the cell, followed by glycerol production either *via* photosynthetic CO_2_ fixation or starch degradation [43, 44]. The produced glycerol serves as an osmoprotectant, to avoid bursting of the cell [42]. The level of intracellular glycerol is proportional to the extracellular salt concentration, reaching values higher than 50% of the dry weight of the cell [45].

This paper aims to optimize the overall glycerol titer extracted from algae in respect to potential future applications, such as the pharmaceutical industry or the production of green chemicals. Different *Dunaliella* strains from Australia were isolated and their potential to accumulate glycerol was investigated. The most promising candidates were applied in a process to optimize parameters that impact glycerol synthesis, such as light, phosphate- and nitrogen limitation, as well as the NaCl-concentration [43, 46, 47]. The optimized conditions were combined to further improve the glycerol titer.

## Materials and Methods

### Algae strains and cultivation

*D. tertiolecta* UTEX 999 was obtained from the Culture Collection of Algae at the Georg August University of Göttingen. The strains #006, #27, #37, #83, # 96, #101 and #127 were isolated from Australian environmental samples obtained in August 2014 by Thomas Brück and coworkers under permission of the Australian government [46]. The purity of the derived unialgal cultures was verified by microscopy (Zeiss AxioLab A.1, Carl Zeiss AG, Oberkochen, Germany).

### Microalgae cultivation

All microalgae strains were cultured in 500 mL Erlenmeyer flasks with a culture volume of 200 mL in New Brunswick Innova 44 series shakers (28°C, 120 rpm) fitted with light emitting diodes (LED) (Future LED GmbH, Berlin, Germany) as described by Woortmann et al. [48]. If not otherwise mentioned, a continuous illumination at a Photosynthetic Photon Flux Density (PPFD) of 100 μmol m^−2^ s^−1^ between 400 nm and 750 nm was applied, to achieve a visible sunlight spectrum (color spectrum AM 1.5 G). Each flask was aeriated with 2% v/v CO_2_ enriched air which was controlled by a DASGIP® MX module (Eppendorf AG, Hamburg, Germany). The microalgae starting with an OD_750_ of 0.1 were cultivated in modified Johnson medium at a pH of 7.5 [49] with 1 M NaCl, if not other specified.

### Phylogenetic characterization of strains

DNA of the strains was extracted by the InnuPrep plant DNA extraction kit (Analytic Jena AG, Jena, Germany, 845-KS-1060050). The strains were identified as *Dunaliella* strains by the amplification of the 18S rDNA using the primers EukA (21F) (AACCTGGTTGATCCTGCCAGT) and EukB (1791R) (GATCCTTCTGCAGGTTCACCTAC) [50]. To further identify the species, the internal transcribed spacer (IST)-sequence as well as the rubisco gene were amplified. Therefore, the ITS Primer ITS3 (GCATCGATGAAGAACGCAGC) and ITS4 (TCCTCCGCTTATTGATATGC) [51] and the rubisco primer rbcLaF (ATGTCACCACAAACAGAGACTAAAGC) and rbcLaR (GTAAAATCAAGTCCACCRCG) [52] were used. The purified amplicons were sequenced by sanger sequencing (Eurofins Genomics GmbH, Ebersberg, Germany) and the obtained sequence were searched against the GenBank database by the BLASTn algorithm [53].

### Glycerol extraction and measurement

After 14 days of cultivation the NaCl concentration was abruptly increased, if not stated otherwise, to 2 M by adding the same volume of 3 M NaCl solution. As a control, instead of 3 M the same volume of a 1 M NaCl solution was added. To measure the accumulated glycerol, the biomass was harvested by centrifugation (2500 g, 5 min). The supernatant was removed. Half of the initial volume of bi-distilled H_2_O was added to the cell pellet. Cells were resuspended and centrifuged at 15,000 g for 5 min. After repeating the previous step, the glycerol content of the extract was determined either using the Glycerin-Assay-Kit (MAK117, Sigma USA) or the Free Glycerol Determination Kit (FG0100, Sigma USA). In addition to the manufacturer’s instructions, for the Free Glycerol Determination Kit (FG0100), sample and reagent volumes was halved. Modified Johnson medium was used as blank.

### Nitrogen- and phosphate limitation

Stress such as nutrient limitation often leads to the production of secondary metabolites [54]. For that reason, the impact of a nitrogen-or phosphate limitation on the glycerol production was investigated. Instead of adding a 3 M NaCl solution to the culture, after 14 days, the culture was centrifuged at 2500 g for 5 min. The old medium was removed and modified Johnson medium either with or without nitrogen and phosphate, respectively, was added to the cells. To provide the cells with sufficient time to adapt to the limited conditions, after 24 h and 78 h also the glycerol concentration after 78 h was measured.

### Analyzing the impact of salt

Since glycerol serves as an osmoprotectant to the culture’s salt concentration, salt influence the glycerol titer [55]. To investigate the impact of the salt concentration, the algae were cultivated at different salt concentration (1 M, 1.5 M and 2 M) and after 14 days the NaCl concentration was raised to 2 M, 3 M and 4 M by adding NaCl solutions in the required molarity. As a control the cultures were diluted with the same volume, but the salt concentration was retained constant.

### Analyzing the impact of light

As the growth of algae is highly related to photosynthesis [30, 31], the impact of light on the growth of the algae, as well as their glycerol production potential, was evaluated. For this purpose, the algae were cultivated with different light intensities and -colors. To analyze the impact of the light intensity, the algae were cultivated with 100, 500, 1000, 1500, and 2000 μmol m^−2^ s^−1^ visible sunlight spectrum (in the following called white light). Additionally, the effect of red and blue light on the algae growth and glycerol concentration was investigated. Thus, 100 and 1000 μmol m^−2^ s^−1^ were adjusted by using only light of the wavelengths 680 nm and 740 nm for red light conditions and 425 nm, 455 nm, and 470 nm for blue light.

### Improved conditions

As a final experiment, the best conditions of the previous investigations were combined to optimize the glycerol titer. In addition to *D. tertiolecta* and #96, the two isolates #27 and #83 were tested. Optimized cultures were cultivated with a light intensity of 500 μmol m^−2^ s^−1^ with 1.5 M NaCl concentration. After 14 days of cultivation, salt was increased to 3 M NaCl. The non-optimized condition was cultivated as mentioned before.

### Nuclear Magnetic Resonance (NMR) spectroscopy

For NMR analysis, glycerol was extracted with ethanol instead of water. Therefore, the biomass was lyophilizated (Zirbus Technology, Bad Grunz, Germany) before glycerol was extracted with GC-grade ethanol. The ethanol extract was treated with activated carbon. After centrifugation at 15,000 g for 5 min, supernatant was lyophilizated again and algae derived glycerol remained. 1H-NMR spectra of algae extracted glycerol were recorded on a Bruker Ascend 400 MHz spectrometer. All spectra are referenced on the proton signal of CDCl3 (7.26 ppm).

### Statistical

At least triplicates were measured for each treatment in the experiment. The results are expressed as mean ± standard deviation in the figures. For improved readability, only mean values are stated in the text. A statistical analysis of differences between tested conditions compared to the control (100 μmol m^−2^ s^−1^, white light, cultivated at 1 M followed by an increase to 2 M) was performed *via* t-test and the level of statistical significance was set to p < 0.05.

## Results

The reference strain, *D. tertiolecta*, was used to investigate whether glycerol is produced intracellularly only or if the cells also secrete glycerol into the medium, as published by Chow et al. [44]. In the present study, glycerol was found only inside of the cell, and nothing was present in the surrounding medium (SI 1). These results are in alignment with Ben-Amotz et al., who described that the major function of glycerol is to maintain the osmotic balance [56]. As no glycerol was found in the medium, only the intracellular glycerol was analyzed in the subsequent experiments.

By 18S-analysis, the seven isolates were identified as *Dunaliella* strains. In addition, the analysis of the ITS sequence confirmed them as *Dunaliella* strains (SI 2). The sequence of the rubisco gene analysis did cause ambiguous results, as the available database for algae was insufficient. Even though all isolates belong to the same genus, they differ in their phenotypical appearance. While some species only show a size of 5 µm, others grow up to a diameter of 15 µm, such as *D. tertiolecta* (Figure 1).

**Figure 1.**
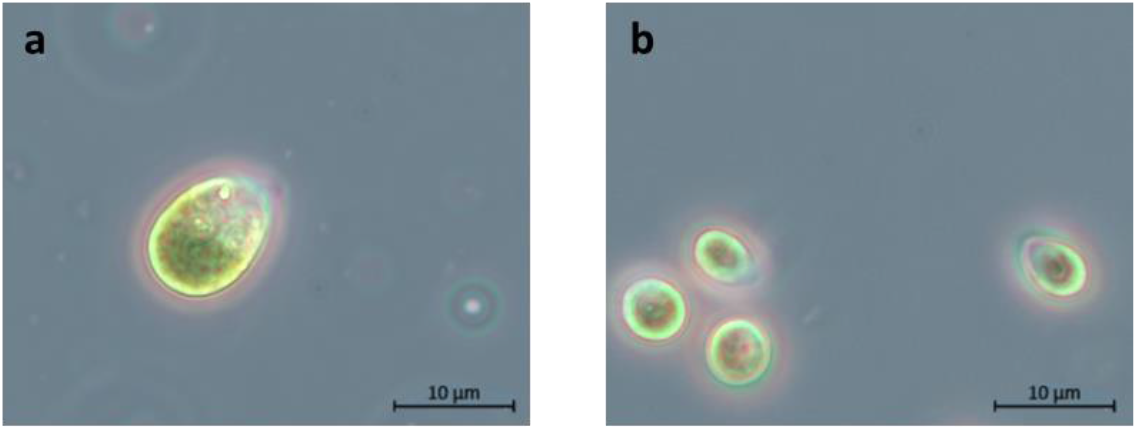
Bright field images (100x) of the morphology of two strains of *Dunaliella*. **a)** *Dunaliella tertiolecta* (UTEX 999). **b)** *Dunaliella* isolate #127, an environmental isolate.

Figure 2 demonstrates that after 14 days of cultivation, the OD_750_ of the different strains ranged from 1.6 to 3.8. In addition, some strains such as *D. tertiolecta* already enter the stationary phase after 8 days, while other strains such as #96 and #37 continue growing linearly by day 14.

**Figure 2.**
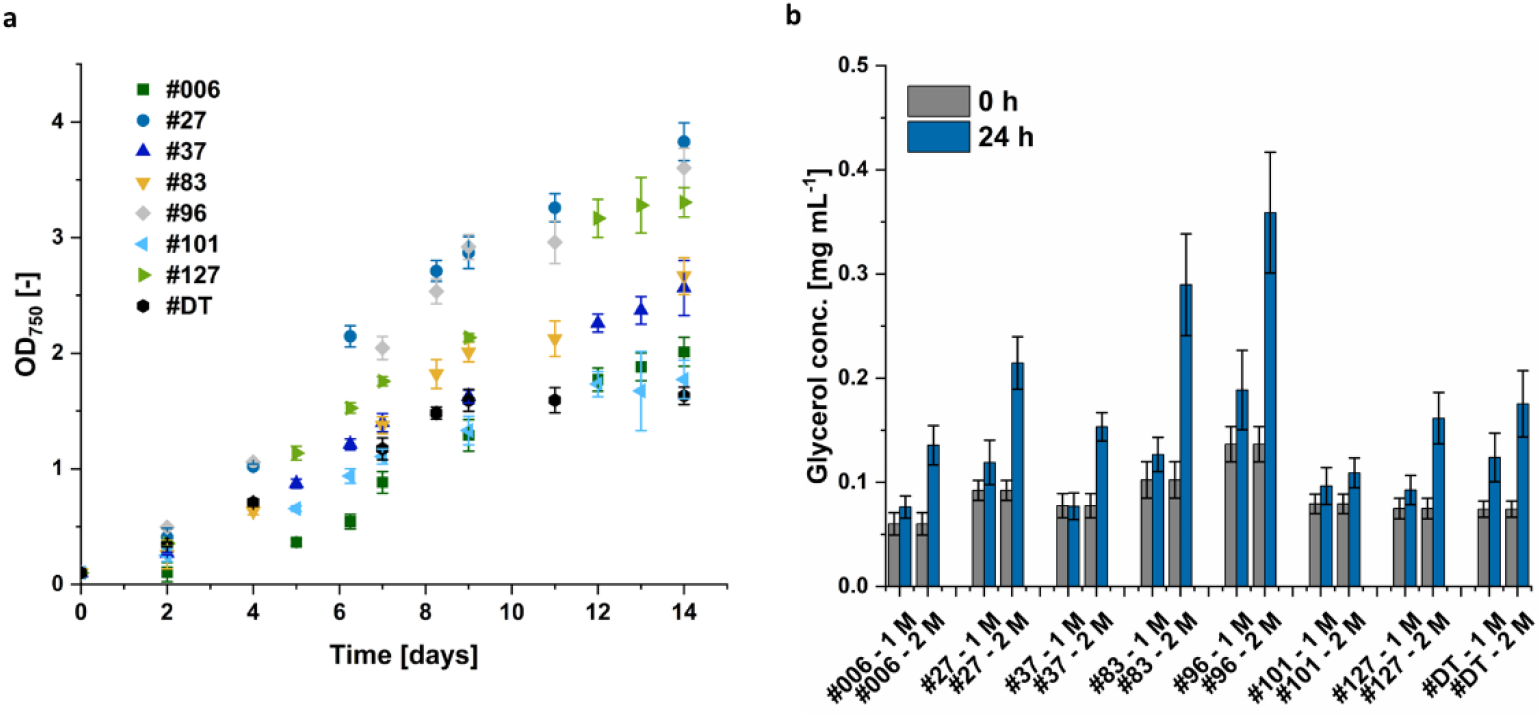
**a)** Growth curve of eight different *Dunaliella* strains. #DT refers to *D. tertiolecta*. #27-#127 are different *Dunaliella* isolates. Algae were cultivated in 1 M NaCl modified Johnson medium with 100 μmol m^−2^ s^−1^ illumination for 14 days. **b)** Glycerol titer of these eight *Dunaliella* strains after 14 days of cultivation (0 h) and 24 h after NaCl concentration was increased to 2 M NaCl or remained constant at 1 M NaCl. After collecting the biomass, glycerol was extracted by adding bi-distilled H_2_O and centrifugation at 15,000 g for 5 min.

The isolates not only varied in size and OD_750_, but also differed in their glycerol production potential. In particular, isolate #96 was identified as a promising candidate for future investigations with a glycerol concentration almost twice as much as *D. tertiolecta* (Figure 2 b). Therefore, this isolate and the reference strain *D. tertiolecta* were chosen for the experimental optimization of the glycerol titer. Afterwards, the optimized conditions were applied to the four most promising candidates: *D. tertiolecta*, #27, #83, and #96.

Initially, the impact of nitrogen- and phosphate limitation was investigated, as limitations are reported to promote the production of secondary metabolites [54]. Therefore, after 14 days of cultivation the medium was removed and new medium with or without nitrogen and phosphate was added. However, the limitations tested in this study neither increased the growth of the algae (data not shown) nor the glycerol concentration after 24 h or 78 h (Figure 3).

**Figure 3.**
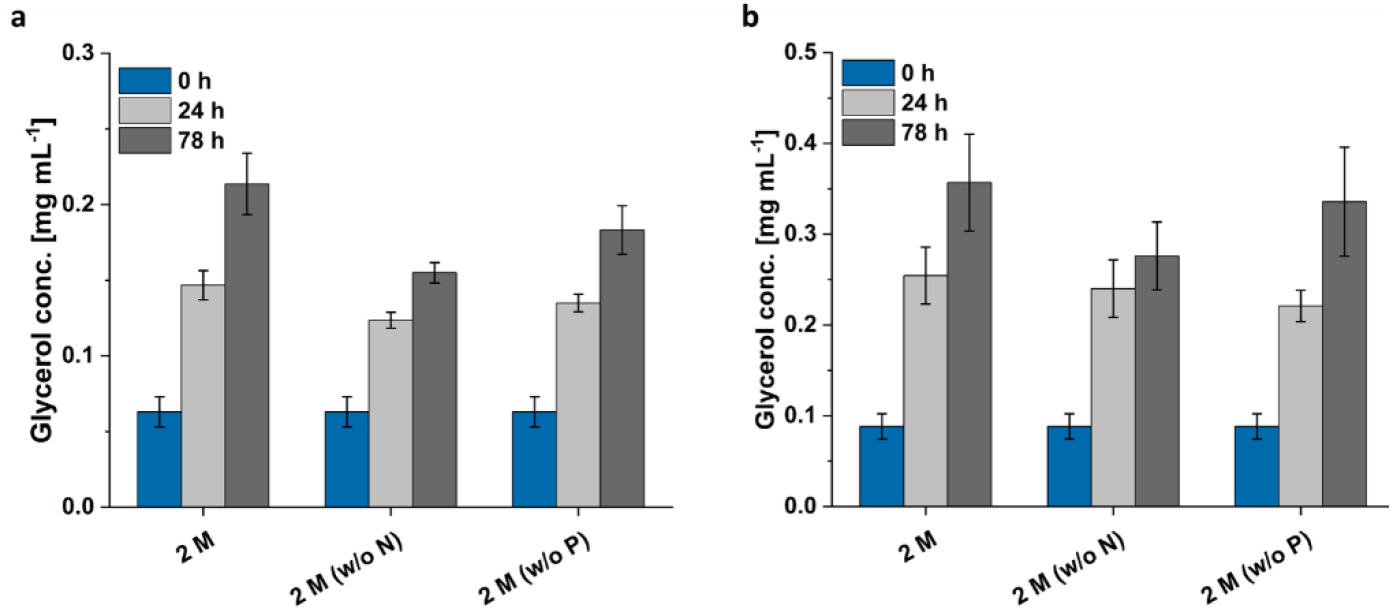
Glycerol concentration of *Dunaliella* strains cultivated 14 days in 1 M NaCl containing full modified Johnson medium (pH=7.5) at 28°C. After 14 days, depleted medium was removed and fresh medium with either 2 M NaCl, 2 M NaCl without nitrogen or 2 M NaCl without phosphate was added. Glycerol concentration was measured at time point 0 h, 24 h and 78 h. **a)** Glycerol concentration of *D. tertiolecta*. **b)** Glycerol concentration of *Dunaliella* isolate #96.

Ben-Amotz et al. analyzed that the accumulated glycerol correlates with the extracellular salt concentration [43, 45]. Accordingly, the algae were cultivated with different NaCl concentration and after two weeks, the NaCl concentration was increased either to 2 M, 3 M, or 4 M. It was decided that the best procedure to study the impact of salt was to cultivate the algae in medium containing 1 M, 1.5 M, and 2 M NaCl, as cells cultivated in 0.5 M NaCl lysed, when the NaCl concentration was suddenly increased to 2 M and 3 M NaCl (data not shown).

Both *Dunaliella* strains grew best in 1 M NaCl. After 14 days *D. tertiolecta* cells reached only an OD_750_ of 1.52 in 1.5 M NaCl and 1.39 in 2 M NaCl compared to 1.66 of the control, cultivated in 1 M NaCl. The deteriorated growth was even more present for isolate #96. The cells grown in 1.5 M, and 2 M NaCl only reached ∼80% and ∼65% of the OD_750_ of the control with a final OD_750_ of 3.0. The #96 growth variations manifested not only by lower biomass formation (Figure 4), changed morphology, but also significant cellular aggregation and clumping as observed in Figure 5. This phenomenon has previously been characterized as the cellular palmella stage [57, 58].

**Figure 4.**
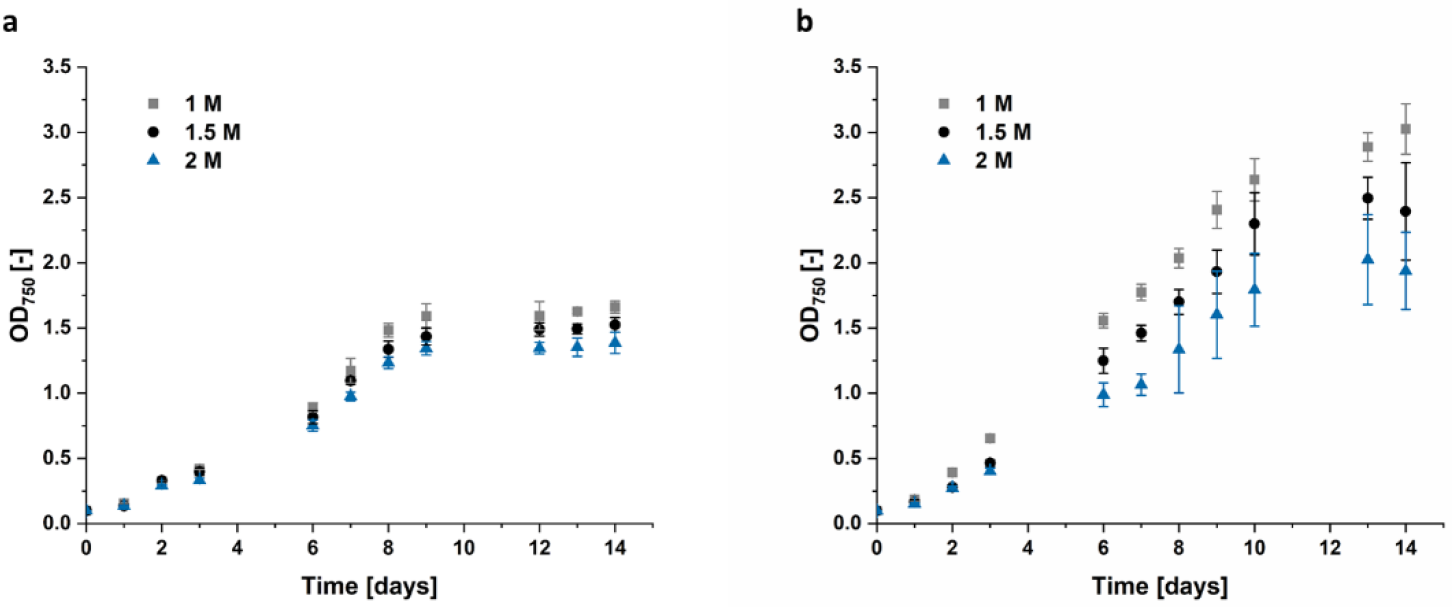
Growth curve of **a)** *D. tertiolecta* and **b)** isolate #96 cultivated in modified Johnson medium (pH=7.5) at 28°C with 100 μmol m^−2^ s^−1^ illumination for 14 days. NaCl concentration of the medium varied between 1 M, 1.5 M, and 2 M.

**Figure 5.**
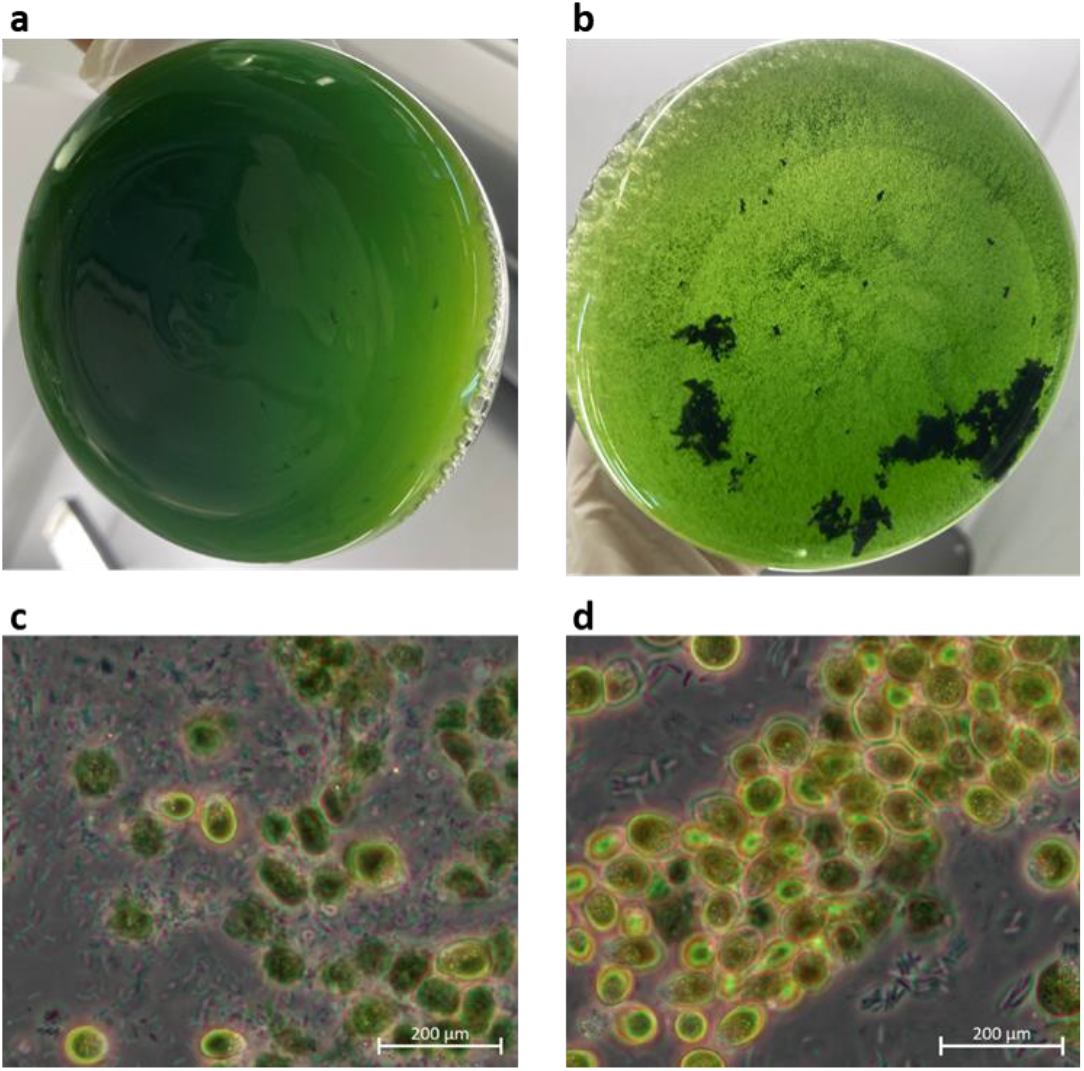
**a)** Bottom view of the flask of *Dunaliella* isolate #96 after one week of cultivation in 1 M NaCl containing modified Johnson medium (pH=7.5) at 28°C. **b)** Bottom view of the flask of isolate #96 after one week of cultivation in 1.5 M NaCl containing modified Johnson medium, demonstrating the forming of accumulated cells. Bright field images (100x) of isolate #96 after hyperosmotic changes. Cells were cultivated with c) 1.5 M or d) 2 M NaCl for 14 days followed by a hyperosmotic change to 4 M NaCl.

*D. tertiolecta*’s highest glycerol titer was achieved when cells were cultured in 1.5 M or 2 M NaCl followed by an increase to 3 M NaCl, but also if cells were cultured in 2 M NaCl with a hyperosmotic change to 4 M NaCl (Figure *5*). The resulting extracts all provided a glycerol concentration of 0.26 mg mL^-1^, increasing the glycerol concentration by 14%. Cells grown in 1 M or 1.5 M did not survive the sudden NaCl increase to 4 M, resulting in lysed cells and almost no extracted glycerol. The cells grown in 1 M NaCl followed by an increase to 3 M NaCl have a high standard deviation which indicates, that some cells already lysed while others could grow at hyperosmotic conditions (Figure 6). This high standard deviation is also visible for isolate #96.

**Figure 6.**
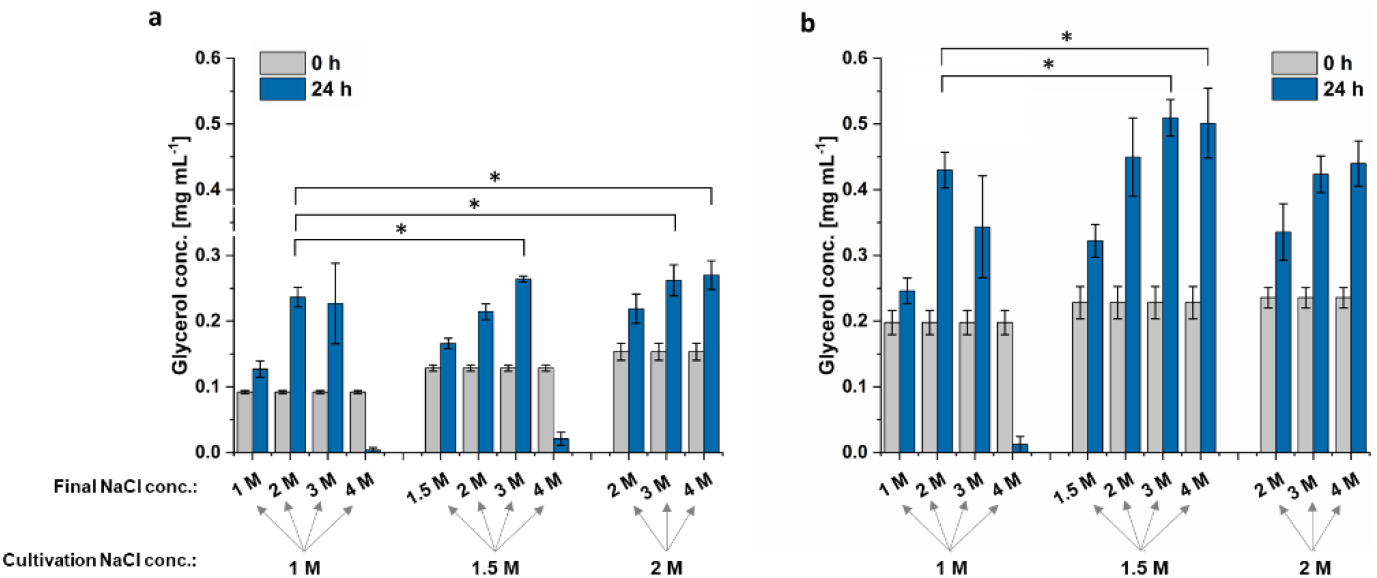
Glycerol concentration of *Dunaliella* strains cultivated 14 days in modified Johnson medium (pH=7.5) at 28°C with 1 M, 1.5 M or 2 M NaCl. After 14 days NaCl concentration was increased to either 2 M, 3 M or 4 M NaCl or kept constant. Glycerol concentration was measured at time point 0 h and 24 h after hyperosmotic changes. **a)** Glycerol concentration of *D. tertiolecta*. **b)** Glycerol concentration of the *Dunaliella* isolate #96. A statistical analysis of differences between tested conditions compared to the control (cultivated at 1 M followed by an increase to 2 M) was performed via t-test and the level of statistical significance (*) was set to p < 0.05.

The highest glycerol concentration extracted from isolate #96 was reached when cells were grown at 1.5 M NaCl followed by an increase to 3 M or 4 M NaCl. These conditions increased the glycerol concentration by 19% to 0.51 mg mL^-1^. Interestingly, although the cells formed clumps when culturing at 1.5 M and 2 M NaCl and had a lower OD_750_ compared to the cells grown in 1 M NaCl, these cells still survived the hyperosmotic change and were able to adapt to the new NaCl concentration by glycerol production. The adapted cells are seen in Figure 5 a and b. When cells were shocked with NaCl from 2 M to 4 M, the cells form more compact clumps. By contrast, when cells were shocked with NaCl from 1.5 M to 4 M, clumps were less compact but cells appear to be less vital. However, even though the formed cell clumps differ in their phenotype after the hyperosmotic change to 4 M, the cells of both conditions survived the sudden salt increase.

The best condition for the NaCl shock appear to be the transition from 1.5 M to 3 M. Even though, other conditions resulted in same increase in glycerol concentration, higher salt load, such as 4 M NaCl, should be avoided, as salt impedes production processes and causes corrosion of steal containing photobioreactor parts at industrial scale [59, 60].

Since the glycerol concentration correlates with the biomass formation, we not only investigated parameters that directly influence glycerol production, but also parameters improving the algae growth pattern. Consequently, the impact of light on algae growth was evaluated. Accordingly, the algae were cultivated with different light intensities (100, 500, 1000, 1500, and 2000 μmol m^−2^ s^−1^) and colors, specifically white, red, and blue light (sun light spectrum AM 1.5 G for white light, 680 nm and 740 nm for red light and 425 nm, 455 nm, and 470 nm for blue light). Interestingly, light intensity but not color had the highest impact on algae growth. With an illumination of white light at 500 μmol m^−2^ s^−1^, the growth of the *D. tertiolecta* increased ∼150%, while at 1000 μmol m^−2^ s^−1^ it increased ∼250% compared to the 100 μmol m^−2^ s^−1^ reference control (Figure 7). Further, the increase to 1500 and 2000 μmol m^−2^ s^−1^ only resulted in slight growth improvements.

**Figure 7.**
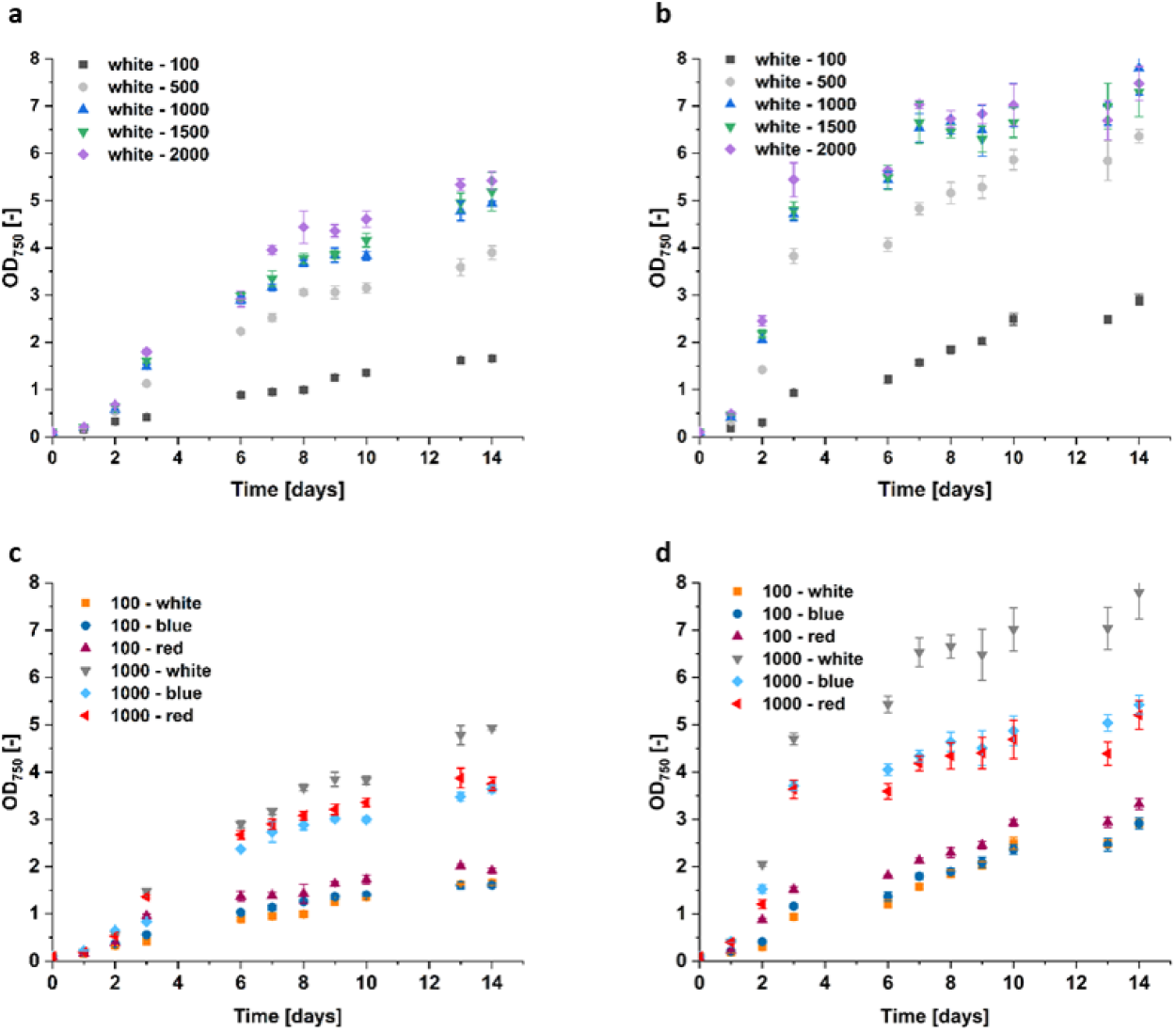
Growth curve of *Dunaliella* grown with different light intensities and colors. **a)** *D. tertiolecta* and **b)** isolate #96 were cultivated in modified Johnson medium (pH=7.5) at 28°C over 14 days with the illumination of white light and varying intensities (100 – 2000 μmol m^−2^ s^−1^). Impact of the illumination of white, red, and blue light with the intensity of 100 and 1000 μmol m^−2^ s^−1^ for **c)** *D. tertiolecta* and **d)** isolate #96 when cultured in modified Johnson medium (pH=7.5) at 28°C over 14 days are shown.

Moreover, the growth of isolate #96 doubled with an increase of white light from 100 to 500 μmol m^−2^ s^−1^, and was even 2.5 times higher when light intensity was increased to 1000 μmol m^−2^ s^−1^. This cell growth could not be improved further by higher light intensities. Similar to cells of #96, grown in 1.5 M and 2 M NaCl, the cells cultivated at light intensities higher than 500 μmol m^−2^ s^−1^ also formed clumps as shown in Figure 6 b. Starting at the fifth day of cultivation at 2000 μmol m^−2^ s^−1^, the higher the light intensity, the earlier the clump formation started. The size of the clumps increased with increasing light intensity as well as with higher cell densities. This indicates that illumination with intensities higher than 500 μmol m^−2^ s^−1^ implies stress for the cells. The red and blue light impacted the growth rate of #96 similar to *D. tertiolecta*., as the resulted OD_750_ was significantly lower compared to the white light at 1000 μmol m^−2^ s^−1^. Interestingly, for both strains at 100 μmol m^−2^ s^−1^, red light resulted in the highest cell growth. However, cells grow best with white light when intensities were raised to 1000 μmol m^−2^ s^−1^.

Figure 8 depicts, that the cultivation at 1000 μmol m^−2^ s^−1^ white light causes higher glycerol titers (*D. tertiolecta*: 0.49 mg mL^-1^, #96: 0.53 mg mL^-1^), compared to red (*D. tertiolecta*: 0.37 mg mL^-1^, #96: 0.23 mg mL^-1^) and blue light (*D. tertiolecta*: 0.40 mg mL^-1^, #96: 0.45 mg mL^-1^), respectively. At 100 μmol m^−2^ s^−1^ there is no significant difference between the light colors, with ∼0.20 mg mL^-1^ for *D. tertiolecta* and 0.29 mg mL^-1^ for #96. The glycerol concentration attains a saturation for light intensities higher than 500 μmol m^−2^ s^−1^, with around 0.45 mg mL^-1^ extracted from *D. tertiolecta* and 0.55 mg mL^-1^ from isolate #96. Thereby, both strains attained double the amount of extracted glycerol compared to the control. Even though higher biomass was achieved with light intensities above 1000 μmol m^−2^ s^−1^ compared to 500 μmol m^−2^ s^−1^, the higher biomass did not result in higher glycerol titer. This again indicated, that light intensities higher than 500 μmol m^−2^ s^−1^ imply stress for the cells. Additionally, it might be assumed, that the glycerol assimilation per cell is limited to a certain amount and decreased with higher light intensities, as in relation to the growth of the cells less glycerol could be extracted. This is currently being investigated in further system biological analysis in our group.

**Figure 8.**
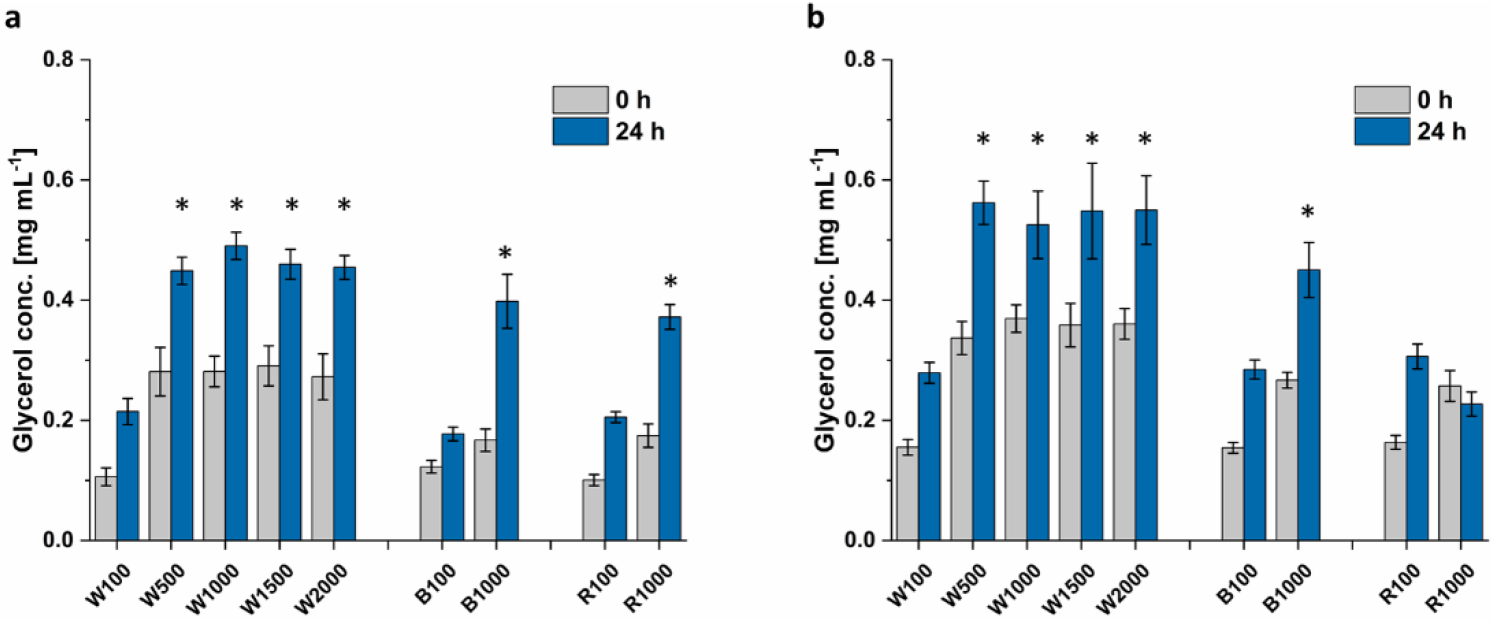
Glycerol concentration of **a)** *D. tertiolecta* and **b)** isolate #96. Algae were cultivated in 1 M modified Johnson medium (pH=7.5) at 28°C with different light intensities and colors. (W=white, B=blue, R=red. The numbers refer to the light intensity in μmol m^−2^ s^−1^). After 14 days of cultivation the NaCl concentration was increased to 2 M. Glycerol concentration was measured at time point 0 h and 24 h after variation of salt concentration. A statistical analysis of differences between tested conditions compared to the control (100 μmol m^−2^ s^−1^, white light) was performed via t-test and the level of statistical significance (*) was set to p < 0.05.

Optimal conditions were identified as white light of 500 µmol m^-2^ s^-1^ and a sudden increase of salt concentration from 1.5 M to 3 M after a growth phase of 14 days. Even though, the optimization was performed with *D. tertiolecta* and isolate #96, the four most promising *Dunaliella* strains according to Figure 1 were cultivated at these optimized conditions, which include beside *D. tertiolecta* and isolate #96 also isolate #27 and #83. The algae growth under optimized conditions was increased by ∼35% for #27, #83, and #96 and even ∼95% for *D. tertiolecta* compared to the non-optimized conditions (Figure 9 a). The highest OD_*750*_ of 4.7 was reached by the small cells of isolate #27, followed by #96 (4.1), #83 (3.3), and *D. tertiolecta* (3.1) when cultured at optimized conditions. Even though, under optimized conditions #27 reached the highest OD_750_, it did not lead to the highest extracted glycerol and resulted in 0.76 mg mL^-1^ only. This concentration is quite similar to the concentration of the extract from the reference strain *D. tertiolecta* (0.79 mg mL^-1^) and isolate #83 (0.74 mg mL^-1^). The extracted glycerol of isolate #96 was 20% higher in comparison to the other strains resulting in a glycerol concentration of 0.94 mg mL^-1^.

**Figure 9.**
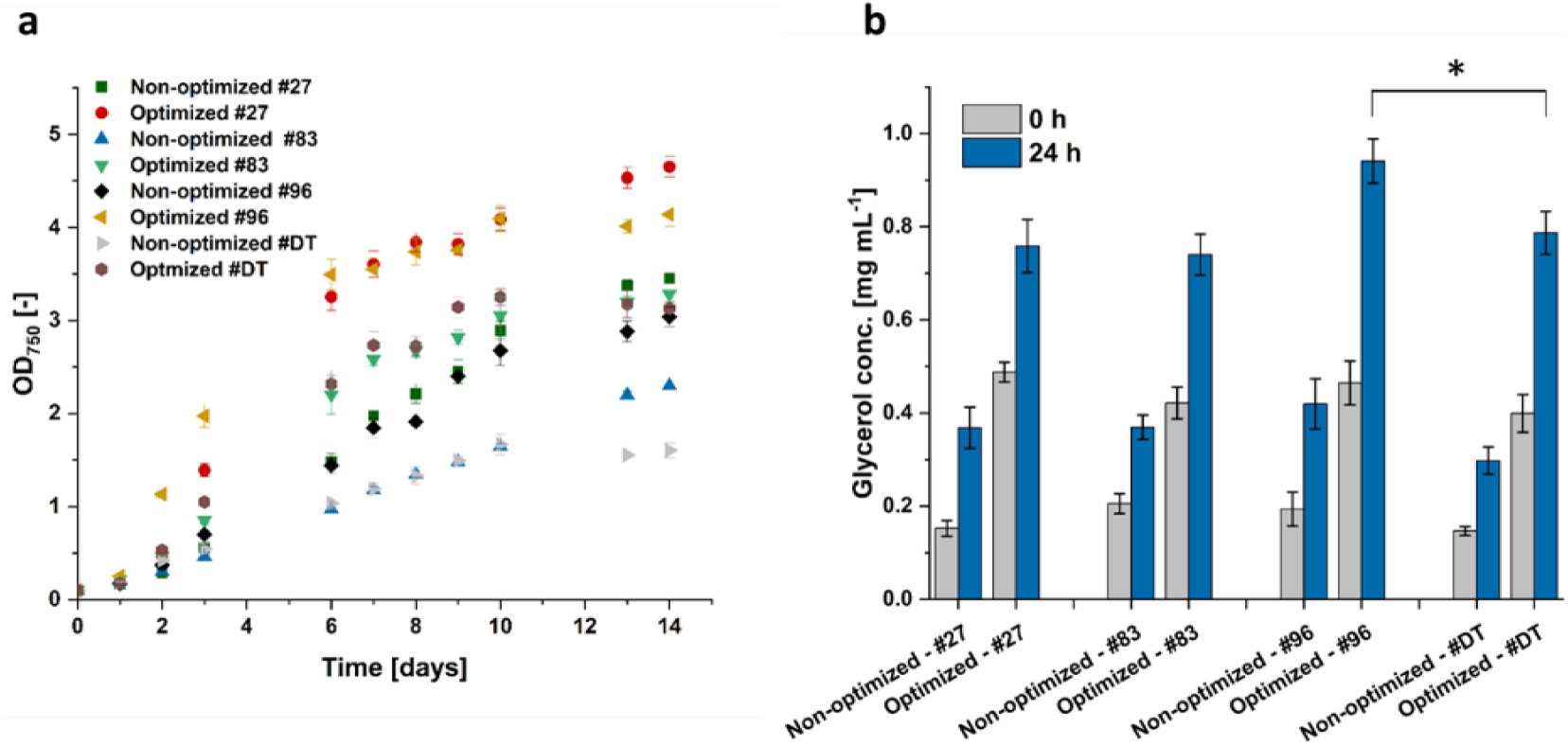
**a)** Growth curve of three *Dunaliella* isolates (#27, #83 and #96) and *D. tertiolecta* (#DT) over 14 days when cultivated under optimized conditions (500 μmol m^−2^ s^−1^ white light and 1.5 M NaCl modified Johnson medium, pH=7.5) or under non-optimized conditions (100 μmol m^−2^ s^−1^ white light and 1 M NaCl modified Johnson medium, pH=7.5) at 28°C. **b)** Glycerol concentration 0 h and 24 h after NaCl concentration was increased to 2 M (non-optimized) or 3 M (optimized). A statistical analysis of differences between optimized conditions of isolates compared to optimized conditions of *D. tertiolecta* was performed via t-test and the level of statistical significance (*) was set to p < 0.05.

Under non-optimized conditions the glycerol concentration extracted from the isolates were higher compared to *D. tertiolecta* which is in alignment with our results shown in Figure 2. With optimized conditions only #96 showed higher glycerol concentration compared to the reference strain *D. tertiolecta*. However, as the conditions were optimized for #96 and *D. tertiolecta*, in future studies the other two isolates, #27 and #83, should be optimized individually to potentially increase their extracted glycerol titer further.

The NMR analysis, shown in Figure 10 indicates that the extract isolated from algae consists of glycerol, when compared to reference H-NMR spectra of glycerol [61]. Additionally, there are some other peaks visible, indicating impurities in the sample. As proteins are almost insoluble in ethanol [62], it is unlikely that proteins are present. The impurities might consist of cell components or carbohydrates, as these are slightly soluble in ethanol [63].

**Figure 10.**
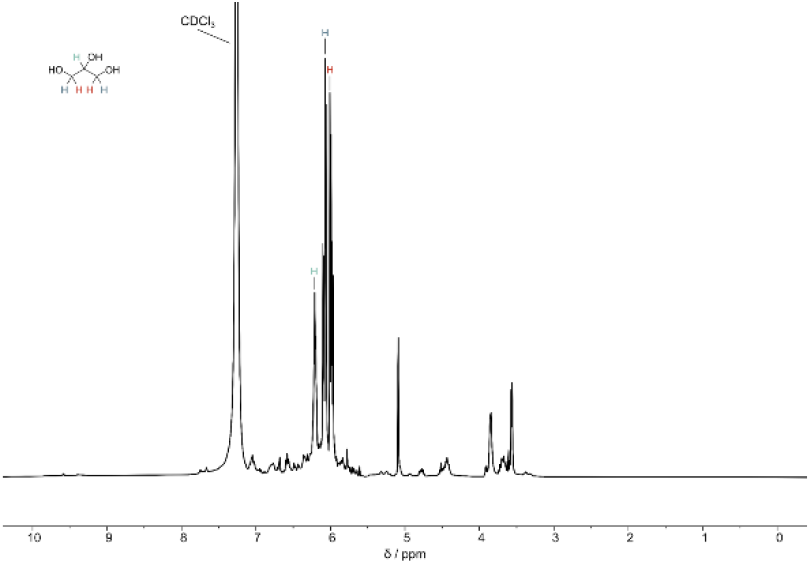
H-NMR spectra of the ethanol extracted glycerol from *Dunaliella tertiolecta* recorded on a Bruker Ascend 400 MHz spectrometer. All spectra are referenced on the proton signal of CDCl3 (7.26 ppm).

## Discussion

The new isolates used in this study, were identified as *Dunaliella*. However, it is challenging to identify the exact species using the 18S, ITS and rubisco gene sequences, as various *Dunaliella* species were suggested with among others 100% identity. This problematic identification is in line with the results from Highfield et al. and Assunção et al., who describe the diversity of the *Dunaliella* strains [64, 65]. Using six different primer pairs, Highfield et al. classified the isolates of their study to one of four sub-clades, but they were not able to determine the exact species. In their opinion, the classification of *Dunaliella* has presented a challenge for many decades and it is necessary, that the scientific community tackles this issue to facilitate the identification of their working strains. The exact identity of a strain would enable targeted selection for specific applications [64].

Even though the isolates all belong to the genus of *Dunaliella*, their cell sizes range from 5 to 15 µm. After 14 days of cultivation their OD_750_ range from 1.6 to 3.8, which is more than double. The difference in size and growth between *Dunaliella* strains has already been observed by Xu et. al. [66], who described a range of 7 µm to 12.5 µm for the size of their isolates. Additionally, the specific growth rate in their study ranged from 0.1 to 0.7 per day at 200 μmol m^−2^ s^−1^. These results highlight the diversity within the *Dunaliella* clades and underscore the requirement for individual investigation of each isolate.

Glycerol was only accumulated inside the cell and was not excreted to the medium, in contrast to the earlier findings of Chow et al. [44]. Due to the differences within the *Dunaliella* genus, it may be assumed, that their strain secretes glycerol, while our strains only accumulate glycerol inside the cell. On the other hand, there is evidence to suggest, that they found extracellular glycerol as a result of high centrifugation at 10,000 g for 20 min. This is confirmed by Harvey et al. [7], who discovered that centrifugal forces above 5,000 g cause a decreased number of intact cells. Therefore, it is feasible that extracellular glycerol was detected because the cells were destroyed, while in fact the intracellular accumulated glycerol was measured.

Nutrient limitations often induce the cell to produce fatty acids or secondary metabolites [54]. It was previously reported that under nitrogen depletion more glycerol could be extracted from *Dunaliella* strains [67]. However, the limitation of nitrogen and phosphate in this study did not cause an increased glycerol titer, but instead a reduction. Even though it is reported, that the synthesis of glycerol from glucose is independent of the availability of nitrogen [42], the insufficient amount of nitrogen in this study leads to a decreased growth and thereby to a reduced glycerol titer as presented in Figure 3. This is in alignment with Kanchan Phadwal et al. [47], who also described that nutrient depletion did not cause increased glycerol accumulation in the cells.

The decreased growth is also visible for the cells grown with insufficient amount of phosphate. Phosphate is required for the glycerol production *via* starch degradation as well as for the photosynthetic glycerol production. To convert starch to glycerol, intermediates such as Glucose-6-phosphate and Fructose-1,6-bisphosphate are required [68]. The glycerol produced through the photosynthetic pathways is also phosphate dependent, as the glycerol is derived from the dihydroxyacetone phosphate of the Calvin cycle [69]. Therefore, the limitations tested in this study, decrease the growth of the algae and thereby the overall glycerol titer.

The cultivation with different salt concentration and hyperosmotic changes to various salt concentration should increase accumulated glycerol. Our results show, that the algae grew best in 1 M NaCl, with less growth in 1.5 M and even less in 2 M as shown in Figure 4. The decreased growth with increasing salt concentration was more evident in isolate #96 than in the reference strain *D. tertiolecta*. While the growth of *D. tertiolecta* was decreased by 8% in 1.5 M NaCl and 17% in 2 M NaCl compared to 1 M NaCl, the growth of #96 was lowered by ∼35% when cultured in 2 M NaCl. Shariati et al. and Borowitzka et al. [43, 55, 70] also discovered varying growth of *Dunaliella* strains at different salinities. Additionally, Xu et al., e.g., describe 0.5 M as the optimal NaCl salinity for one strain, while other strains grow better in 1.5 M [66, 71]. This again highlights the differences between the *Dunaliella* strains and the need to examine each strain individually. The cultivation of isolate #96 at a higher salt concentration forced the cells to accumulate and form clumps with each other. Wei et al. and Borowitzka et al. already characterized these clumps as a palmella stage of the cells [57, 58]. The authors discovered, that cells in the palmella stage lose their eyespot and flagella, and become more circular. Additionally, they reported, that the cells excrete a slime layer, which allows them to divide repeatedly and to form aggregation of green cells [58]. Algae enter the palmella stage, when they are exposed to extreme conditions, such as a decrease or increase in salinity or high light intensities [58]. In this study, the higher the salt concentration, the earlier the cells entered the palmella stage and the bigger the clumps (SI 2). This confirms the findings from Montoya et al. [72], who observed increasing salt concentration as a catalyst for the palmella structure formation of *Dunaliella* cells.

Furthermore, our data indicate that extreme NaCl increases such as 1 M to 4 M NaCl lead to cell-bursting. This is in alignment with Avron and Ben-Amotz [68, 73], who discovered that *Dunaliella* cells are able to physically withstand three-to four-fold increases in salt concentration, while further hyperosmotic stress leads to bursting of the cells. To avoid cell-bursting, a gradual increase of the NaCl concentration may help the cells better adapt and should be investigated in future experiments.

The cells of isolate #96 entered the palmella stage, when they were cultured at 1.5 M and 2 M NaCl. Even though the palmella stage is a stress signal, they were able to survive the hyperosmotic change to 4 M NaCl. It is conceivable, that the palmella stage enabled the survival because the cells on the outside of clumps protected the cells on the inside.

The highest glycerol concentration for both strains was obtained when cells were cultured at 1.5 M NaCl followed by doubling of NaCl concentration. Extracts from *D. tertiolecta* also resulted in the same glycerol concentration when cells were cultured in 2 M NaCl followed by a hyperosmotic change to 3 M or 4 M NaCl. However, these two conditions did not lead to the highest extracted glycerol concentration from isolate #96, but resulted in a concentration in the range of the control. In contrast, isolate #96 cultivation in 1.5 M NaCl followed by an increase to 4 M NaCl resulted in higher glycerol concentration compared to the control, while this condition led to cell-bursting of *D. tertiolecta* cells. This again indicates that cells behave differently to hyperosmotic changes and highlights the necessity to investigate each strain separately. Since the glycerol concentration for both strains was improved, when cells were cultured with 1.5 M NaCl followed by a hyperosmotic change to 3 M NaCl, this condition was chosen to be the best condition.

By investigating the light color and intensity we aimed to improve the growth pattern of the algae. Our results show that the growth of the algae increased with increasing light intensity. This phenomenon was detectable up to an illumination of 1000 μmol m^−2^ s^−1^. A further increase to 1500 or 2000 μmol m^−2^ s^−1^ did not lead to increased growth.

Illumination with an intensity higher than 1000 μmol m^−2^ s^−1^ may cause excitation of the photosystems, which leads to the formation of reactive oxygen species as a by-product. Reactive oxygen species cause irreversible photo-oxidative damage if not corrected by the cell [74], resulting in stunted growth. Interestingly, the results of Sui et al. show, that the growth of their *D. salina* could only be increased up to an intensity of 200 μmol m^−2^ s^−1^, assuming that different *Dunaliella* strains adapted to different requirements of their environmental habitat.

Algae have two photosystems (PS) which are responsible for the photosynthetic fixation of CO_2_ and are specialized to different sections of the light spectrum. The main function of PS I is to form NADPH, while PS II hydrolyses water and synthesizes ATP. In comparison to PS II, which is primarily activated by blue light, PS I is more sensitive to (far-) red light [75]. In our experiments the algae grew best with red light at low light intensity (100 μmol m^−2^ s^−1^), which evidences that PS I might play a bigger role in the photosynthetic activity at low light intensities. However, this requires more detailed investigation. Our results also align with the work of Zhao et al. [76] who discovered that red LED produced the highest cell number of *Chlorella*.

It is well-known, that blue and red light promote optimal algae and plant cultivation because of the corresponding peaks in the absorption spectrum [77, 78]. In our experiments, however, blue illumination resulted in the least biomass, independent of light intensity. Although some plants exhibit improved photosynthesis under blue than under red light [79], Luimistra et al. [80] reported, that cyanobacteria grown under blue light display intracellular changes, that cause a redistribution of energy flow between the two photosystems. This indicates, that blue light may create an imbalance between the two photosystems and thereby hinder the cell growth. This also may be the reason for the slower growth of our *Dunaliella* strains.

Compared to low light intensity, algae grown at higher light intensity (1000 μmol m^−2^ s^−1^) resulted in the highest biomass when illuminated with white light. This suggests, that the composition of the light impacts the photosynthetic activity. Red or blue light on its own may cause an imbalance in the stimulation of the two photosystems and in the electron chain [81]. This imbalance may produce reactive oxygen species and photo oxidative damage, both of which are harmful to the organism [75, 82].

The highest biomass production with white light illumination indicates, that algae improved their photosystems to optimally function under white, sun-like light conditions [78]. As the white light spectrum is preferred, potential large-scale production may be realized in open ponds with sunlight exposure. Although algae glycerol production has not reached industrial scale, *Dunaliella salina* is already cultivated in an open pond system for β-carotene production [40]. In that respect, glycerol production can be realized in a similar process.

Xu et al. and Harvey et al. [40, 66] reported, that their *Dunaliella* strains turned red and produced β-carotene when cultured at high intensities of light or at red light. However, when cultured at light intensities higher than 500 μmol m^−2^ s^−^1, our strains remained green and did not turn red. Additionally, instead of β-carotene production, isolate #96 entered the palmella stage. As already mentioned, algae enter the palmella stage when they are exposed to extreme conditions [57, 58]. This indicates that the algae were stressed by light intensities higher than 500 μmol m^−2^ s^−1^ and consequently did not increase the overall glycerol titer, although they produced more biomass. Moreover, the cumulative data indicates, that *Dunaliella* species either generate pigments or glycerol as a stress response.

Since illumination with 500, 1000, 1500 and 2000 μmol m^−2^ s^−1^ all resulted in approximately the same glycerol concentration, white light with an intensity of 500 μmol m^−2^ s^−1^ was identified as optimal illumination to avoid photo oxidative stress reaction in the cells. If algae-based glycerol production was to be operated on a large scale, the intensity of 500 μmol m^−2^ s^−1^ would reduce cooling needs because it consumes less energy and produces less heat.

Finally, the algae were cultivated under optimized conditions. This implies the cultivation at white light with an intensity of 500 μmol m^−2^ s^−1^ in 1.5 M NaCl, followed by an increase to 3 M NaCl after 14 days. These conditions were applied to the already investigated strains *D. tertiolecta* and isolate #96, as well as to isolates #27 and #83, which were also promising candidates according to Figure 2.

With these optimized conditions only #96 showed higher glycerol concentration compared to the reference strain *D. tertiolecta*. As isolate #27 with the highest OD_750_ did not lead to the highest glycerol concentration, the growth pattern of the algae might be decoupled from the glycerol production. To further analyze the glycerol accumulation potential of each strain individually, the glycerol concentration per cell was calculated (Figure 11).

**Figure 11.**
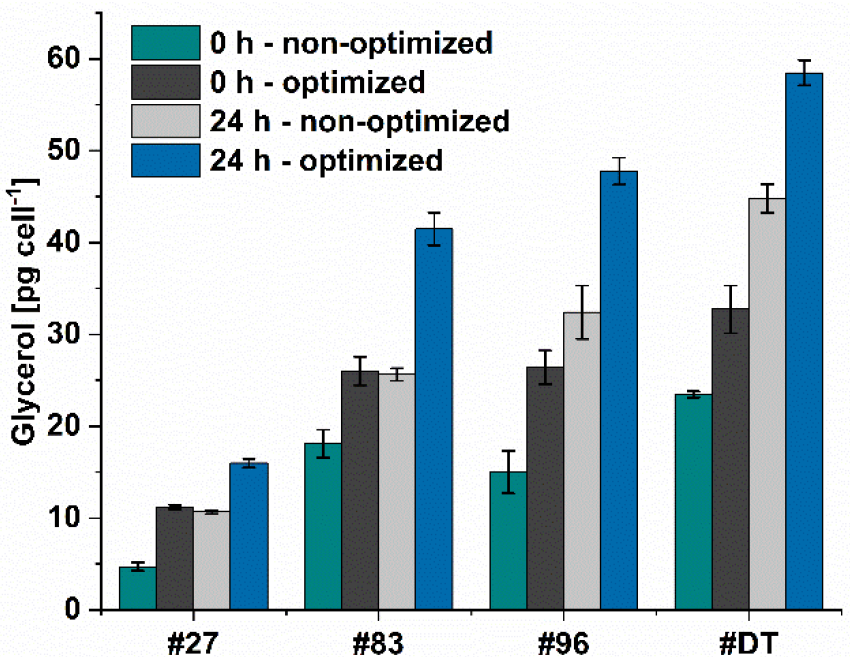
Glycerol concentration per cell for four different *Dunaliella* strains. #DT refers to *D. tertiolecta*, #27-#96 are different *Dunaliella* isolates. Algae were cultivated under optimized conditions (500 μmol m^−2^ s^−1^ white light and 1.5 M NaCl modified Johnson medium, pH=7.5) or non-optimized conditions (100 μmol m^−2^ s^−1^ white light and 1 M NaCl modified Johnson medium, pH=7.5) at 28°C. After 14 days of cultivation NaCl concentration was increased to 2 M (non-optimized) or 3 M (optimized) and glycerol was extracted at time points 0h and 24 h and the amount of glycerol per cell was calculated for these time points.

As for the hyperosmotic change the NaCl concentration was doubled, it was excepted, that the intracellularly accumulated glycerol should double as well. However, the doubled amount of intracellularly accumulated glycerol was only seen for the non-optimized conditions (∼1.94-fold), while for the optimized conditions the intracellularly accumulated glycerol was increase by 1.65 only. This indicates that the amount of glycerol within the cell is limited, which also explains the results from Figure 8, where higher light leads to better growth but not to higher glycerol titer.

Since the isolate #27 is the smallest of the four examined *Dunaliella* strains in this study, expectantly the glycerol amount per cell is the lowest (max. 16 pg cell^-1^) and achieves approximately only one fourth of *D. tertiolecta’s* accumulated glycerol (max. 59 pg cell^-1^). The accumulated glycerol of isolates #83 and #96, the size of which is between isolate #27 and *D. tertiolecta*, reached 71% and 75% compared to *D. tertiolecta*, respectively. These results are in alignment with Xu et al. [66], who’s algae accumulated between 25 and 200 pg glycerol per cell when cultured at 1.5 M. However, the alga that accumulated up to 200 pg cell^-1^ showed the slowest growth rate in their study and thus, does not provide a suitable alternative to our isolate. Interestingly, the cells of isolate #27 and #83 accumulate equal amount of glycerol at 0 h and optimized conditions (1.5 M), as well as after 24 h of non-optimized conditions (2 M). In contrast, *D. tertiolecta* and isolate #96 accumulate higher amounts of glycerol at 2 M NaCl (24 h – non-optimized), as assumed. These differences indicate the need for system biological analysis of the algae to further understand the mechanism how *Dunaliella* cells adapt to hyperosmotic changes at cell level.

## Conclusion

Seven algae strains from different environmental sites were genetically identified as *Dunaliella* strains. Their capacity to generate glycerol compared to the reference strain *Dunaliella tertiolecta* was examined in this study. Experimental variation of salt concentration and light intensities showed potential to improve overall glycerol titer from the reference *D. tertiolecta* and the proprietary isolates #27, #83, and #96. Optimal conditions were identified with white light of 500 µmol m^-2^ s^-1^ and a sudden increase of salt concentration from 1.5 M to 3 M after a growth phase of 14 days. With these improved conditions, the glycerol concentration for *D. tertiolecta* could be doubled to 0.79 mg mL^-1^ compared to former condition. Glycerol titer in extracts from isolate #96 could be further improved to 0.94 mg mL^-1^. Consequently, the overall glycerol titer could be increased 2.2-fold (isolate #96) and 2.6-fold (*D. tertiolecta*). The improved glycerol titer is beneficial for the potential industrial production of glycerol from *Dunaliella* which allows the cultivation on wasteland and without competition to the food production. This study indicates, that glycerol production for green chemicals has industrial potential, if the correct cultivation conditions and strain selection is applied. Moreover, the physiological and phenotypical differences of *Dunaliella* strains apparent not only in this study suggests, that the *Dunaliella* clade requires a more defined genetic resolution to allow a distinct assignment of working strains under study. Further, the biochemical bases and metabolic networks leading to glycerol formation under differently timed physiological cues should be examined using advanced systems biology tools. This would allow the identification of metabolic bottlenecks in glycerol production, which could either be addressed by genetic engineering or process design intervention. Combining the synergistic power of genetic-with process optimization, may lead towards a techno economically and ecologically viable glycerol production process. In this context, algae-based glycerol synthetized from atmospheric CO_2_ can be converted to ‘green chemicals’ to replace several fossil-based chemicals, such polyacrylonitrile, which among others serves as a precursor for sustainable carbon fibers.

## Acknowledgments and funding information

This project, clean carbon, was funded by the German Federal Ministry for Economic Affairs and Climate Action (Grant No.: 20E1902). TB gratefully acknowledges funding of the Werner Siemens foundation for establishing the research field of Synthetic Biotechnology at the Technical University of Munich.

## Data Availability statement

All data generated or analyzed during this study are included in this published article. However, if detailed values, e.g., if the mean is presented only, are desired, these data are available from the corresponding author on reasonable request.

