## Supplementary for "Optimization of the glycerol production from *Dunaliella tertiolecta* and *Dunaliella* isolates": Supplementary.docx

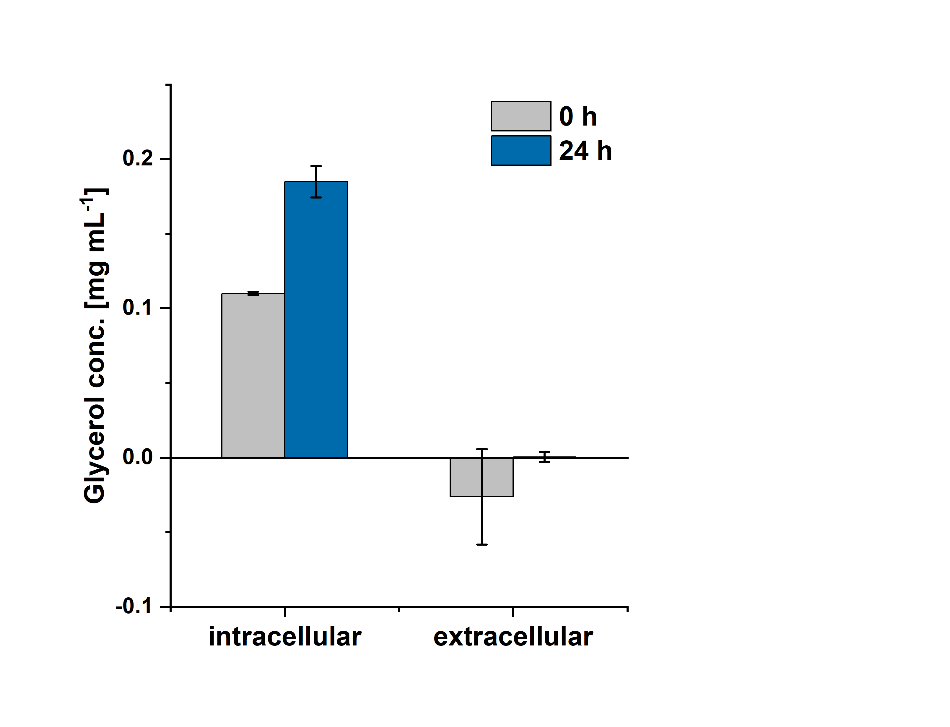

**SI 1** Glycerol concentration of intracellularly and extracellularly derived glycerol, extracted from *D. tertiolecta* after cultivation at 28°C in 1 M NaCl containing modified Johnson medium (pH=7.5). After 14 days, the NaCl concentration was increased to 2 M and glycerol concentration was measured 0 h and 24 h after hyperosmotic change.

**SI 2** FASTA sequence of the 18S Gene sequence of all there tested *Dunaliella* strains

> #006_18S

GGAAGCTGCTAAGATTAAGCCATGCATGTCTAAGTATAAACTGCTTATACTGTGAAACTGCGAATGGCTCATTAAATCAGTTATAGTTTATTTGATGGTACCTTTACTCGGATAACCGTAGTAATTCTAGAGCTAATACGTGCGTAAATCCCGACTTCTGGAAGGGACGTATTTATTAGATAAAAGGCCAGCCGGGCTTGCCCGACTCTTGGCGAATCATGATAACTTCACGAATCGCACGGCTTCGGCCGGCGATGTTTCATTCAAATTTCTGCCCTATCAACTTTCGATGGTAGGATAGAGGCCTACCATGGTGGTAACGGGTGACGGAGGATTAGGGTTCGATTCCGGAGAGGGAGCCTGAGAAACGGCTACCACATCCAAGGAAGGCAGCAGGCGCGCAAATTACCCAATCCCAACACGGGGAGGTAGTGACAATAAATAACAATACCGGGCATTTTTGTCTGGTAATTGGAATGAGTACAATCTAAATCCCTTAACGAGTATCCATTGGAGGGCAAGTCTGGTGCCAGCAGCCGCGGTAATTCCAGCTCCAATAGCGTATATTTAAGTTGTTGCAGTTAAAAAGCTCGTAGTTGGATTTCGGGTGGGTTGTAGCGGTCAGCCTTTGGTTAGTACTGCTACGGCCTACCTTTCTGCCGGGGACGAGCTCCTGGGCTTAACTGTCTGGGACTCGGAATCGGCGAGGTTACTTTGAGTAAATTAGAGTGTTCAAAGCAAGCATACGCTCTGAATACATTAGCATGGAATAACACGATAGGACTCTGGCTTATCTTGTTGGTCTGTAAGACCGGAGTAATGATTAAGAGGGACAGTCGGGGGCATTCGTATTTCATTGTCAGAGGTGAAATTCTTGGATTTATGAAAGACGAACTTCTGCGAAAGCATTTGCCAAGGATGTTTTCATTAATCAAGAACGAAAGTTGGGGGCTCGAAGACGATTAGATACCGTCGTAGTCTCAACCATAAACGATGCCGACTAGGGATTGGCAGGTGTTTCGTTAATGACCCTGCCAGCACCTTATTGAGAAATCAAAGTTTTTGGGTTCCGGGGGGAGTATGGTCGCAGGCTGAAACTTAAAGCGAATTGACGAACGGCCCCCCAGGGGTGGAGCCTGCGGCTTAATTTGACTCAACCCGGAAAACTTACCAGGTCAGACCGGGGAGGATTGACGATTGAAAGCTCTTTCTTGATTTGGGGGGGGGTGCAGGGCCGTCTTATTGGGGGTTGCCTGGCAGGTGAATCCGGAACAAACGGAACCCCACCCGCAAAAAAGCCCCTCTTCCCCGGCGGCCCCCGATTTTTTAAGGGCAAGGGCGTTTTCCAAGGGAAGTGGCGGAAAACAAGGGGTGTAGGCCCTTATTGAGGGGGCCCCGCGCCCCCAACGATATTTTAAGACAACCCTGTGGAAACGGGGCGGGAATTTTA

> #27_18S

TGAGCCTGCATGTCTAAGTATAAACTGCTTATACTGTGAAACTGCGAATGGCTCATTAAATCAGTTATAGTTTATTTGATGGTACCTTTACTCGGATAACCGTAGTAATTCTAGAGCTAATACGTGCGTAAATCCCGACTTCTGGAAGGGACGTATTTATTAGATAAAAGGCCAGCCGGGCTTGCCCGACTCTTGGCGAATCATGATAACTTCACGAATCGCACGGCTTCGTGCCGGCGATGTTTCATTCAAATTTCTGCCCTATCAACTTTCGATGGTAGGATAGAGGCCTACCATGGTGGTAACGGGTGACGGAGGATTAGGGTTCGATTCCGGAGAGGGAGCCTGAGAAACGGCTACCACATCCAAGGAAGGCAGCAGGCGCGCAAATTACCCAATCCCAACACGGGGAGGTAGTGACAATAAATAACAATACCGGGCATTTTTGTCTGGTAATTGGAATGAGTACAATCTAAATCCCTTAACGAGTATCCATTGGAGGGCAAGTCTGGTGCCAGCAGCCGCGGTAATTCCAGCTCCAATAGCGTATATTTAAGTTGTTGCAGTTAAAAAGCTCGTAGTTGGATTTCGGGTGGGTTGTAGCGGTCAGCCTTTGGTGAGTACTGCTACGGCCCACCTTTCTGCCGGGGACGTGCTCCTGGGCTTAACTGTCCGGGACACGGAATCGGCGAGGTTACTTTGAGTAAATTAGAGTGTTCAAAGCAAGCCTACGCTCTGAATACATTAGCATGGAATAACACGATAGGACTCTGGCTTATCTTGTTGGTCTGTAAGACCGGAGTAATGATTAAGAGGGACAGTCGGGGGCATTCGTATTTCATTGTCAGAGGTGAAATTCTTGGGATTTATGAAAGACGAACTTCTGCGAAAGCATTTGCCAAGGATGTTTTCATTAATTCAAGAACGAAAGTTGGGGGCTCGAAGACGATTAGATACCGTCGTAGTCTCAACCATAAACGATGCCGACTAGGGATTGGCAGGTGTTTCGTTGATGACCCTGCCAGCACCTTATGAGAAAATCAAAGTTTTTGGGTTCCGGGGGGAATATGGTCCCAAGGGTTGAAACTTTAAAGGGAATTTGACGGAAGGGGCCCCCCCCGGGGGGGGAACCCGGGGGGTTAATTTGTTCCCCAACCCGGGGAAAAATTTACACGGCCCAAAAAGGGGAGAATTGAAAAATTGAAAATCTTTTTTTTTTTTTG

> #37_18S

GCGCGAATCTGTCTCAAGATTAAGCCATGCATGTCTAAGTATAAACTGCTTATACTGTGAAACTGCGAATGGCTCATTAAATCAGTTATAGTTTATTTGATGGTACCTTTACTCGGATAACCGTAGTAATTCTAGAGCTAATACGTGCGTAAATCCCGACTTCTGGAAGGGACGTATTTATTAGATAAAAGGCCAGCCGGGCTTGCCCGACTCTTGGCGAATCATGATAACTTCACGAATCGCACGGCTTCGTGCCGGCGATGTTTCATTCAAATTTCTGCCCTATCAACTTTCGATGGTAGGATAGAGGCCTACCATGGTGGTAACGGGTGACGGAGGATTAGGGTTCGATTCCGGAGAGGGAGCCTGAGAAACGGCTACCACATCCAAGGAAGGCACCTGGCGCGCAAATTACCCGATCCCAACACGGGGAGGTAGTGACAATAAATAACAATACCGGGCATTTTTGTCTGGTAATTGGAATGAGTACAATCTAAATCCCTTAACGAGTATCCGTTGTAGGGCTAGTCTGGTGCCTGCAGCCGCGGTAGATCCAGCTCCAATAGCGTATATTTAAGTTGTTGCAGTTGAAAAGCTCCGTAGATGGATTTCGGGTGAGTTGTACGAACAGACTTTGCCGAGTACTGCTAACGTGACTACCTTTCATGCCGAGAACGTGACTCCTGGGCATAAACTGTCACGGTAACCACCGTAATGCGGACGAAGGTATATCTTTTGAAGTAATAGTTACTCCGTGCATCTTTA

> #83_18S

CGGCGAATGGCTCATTAAATCAGTTATAGTTTATTTGATGGTACCTTTACTCGGATAACCGTAGTAATTCTAGAGCTAATACGTGCGTAAATCCCGACTTCTGGAAGGGACGTATTTATTAGATAAAAGGCCAGCCGGGCTTGCCCGACTCTTGGCGAATCATGATAACTTCACGAATCGCACGGCTTCGTGCCGGCGATGTTTCATTCAAATTTCTGCCCTATCAACTTTCGATGGTAGGATAGAGGCCTACCATGGTGGTAACGGGTGACGGAGGATTAGGGTTCGATTCCGGAGAGGGAGCCTGAGAAACGGCTACCACATCCAAGGAAGGCAGCAGGCGCGCAAATTACCCAATCCCAACACGGGGAGGTAGTGACAATAAATAACAATACCGGGCATTTTTGTCTGGTAATTGGAATGAGTACAATCTAAATCCCTTAACGAGTATCCATTGGAGGGCAAGTCTGGTGCCAGCAGCCGCGGTAATTCCAGCTCCAATAGCGTATATTTAAGTTGTTGCAGTTAAAAAGCTCGTAGTTGGATTTCGGATGGGTTGTAGCGGTCAGCCTTTGGTGAGTACTGCTACGGCCCATCTTTCTGCCGGGGACGTGCTCCTGGGCTTAACTGTCCGGGACACGGAATCGGCGAGGTTACTTTGAGTAAATTAGAGTGTTCAAAGCAAGCCTACGCTCTGAATACATTAGCATGGAATAACACGATAGGACTCTGGCTTATCTTGTTGGTCTGTAAGACCGGAGTAATGATTAAGAGGGACAGTCGGGGGCATTCGTATTTCATTGTCAGAGGTGAAATTCTTGGGATTTATGAAAGACGAACTTCTGCGAAAGCATTTGCCAAGGATGTTTTCATTAATCAAGAACGAAAGTTGGGGGCTCGAAGACGATTAGATACCGTCGTAGTCTCAACCATAAACGATGCCAACTAGGGATTGGCAGGGGTTTTCGTTGATGACCCTGCCGCACCTTATGAGAAATCAAAGTTTTGGGTTCCGGGGGGGGATAATGGGCGCCAAGGCTGAAACTTTAAGGGAATTTGACGAGAGGGCCCACCCGGCGGGGGAGCCGCGGGTTATTTTGATCCACACGGGAAAATTTCAGGCCGCCCCCGGGGGGGATGTTAAAATTGAAACCTTCTGTTTGTGGGGGGGGGGGGGGGGGGGCCCCTCCTTCTGTGGTGGGGGTGTGTTTGGTGTTTTTGTGTAGAAAAAGAAAACCCCCCGCCGAGAAAAACCCCCCCCCCCCGGGGGCCGGCCTTTTATAGGA

> #96_18S

TCTAAGTATAAACTGCTTATACTGTGAAACTGCGAATGGCTCATTAAATCAGTTATAGTTTATTTGATGGTACCTTTACTCGGATAACCGTAGTAATTCTAGAGCTAATACGTGCGTAAATCCCGACTTCTGGAAGGGACGTATTTATTAGATAAAAGGCCAGCCGGGCTTGCCCGACTCTTGGCGAATCATGATAACTTCACGAATCGCACGGCTTCGTGCCGGCGATGTTTCATTCAAATTTCTGCCCTATCAACTTTCGATGGTAGGATAGAGGCCTACCATGGTGGTAACGGGTGACGGAGGATTAGGGTTCGATTCCGGAGAGGGAGCCTGAGAAACGGCTACCACATCCAAGGAAGGCAGCAGGCGCGCAAATTACCCAATCCCAACACGGGGAGGTAGTGACAATAAATAACAATACCGGGCATTTTTGTCTGGTAATTGGAATGAGTACAATCTAAATCCCTTAACGAGTATCCATTGGAGGGCAAGTCTGGTGCCAGCAGCCGCGGTAATTCCAGCTCCAATAGCGTATATTTAAGTTGTTGCAGTTAAAAAGCTCGTAGTTGGATTTCGGATGGGTTGTAGCGGTCAGCCTTTGGTGAGTACTGCTACGGCCCATCTTTCTGCCGGGGACGTGCTCCTGGGCTTAACTGTCCGGGACACGGAATCGGCGAGGTTACTTTGAGTAAATTAGAGTGTTCAAAGCAAGCCTACGCTCTGAATACATTAGCATGGAATAACACGATAGGACTCTGGCTTATCTTGTTGGTCTGTAAGACCGGAGTAATGATTAAGAGGGACAGTCGGGGGCATTCGTATTTCATTGTCAGAGGTGAAATTCTTGGATTTATGAAAGACGAACTTCTGCGAAAGCATTTGCCAAGGATGTTTTCATTAATCAAGAACGAAAGTTGGGGGCTCGAAGACGATTAGATACCGTCGTAGTCTCAACCA

TAAACGATGCCGACTAGGGATTGGCAGGTGTTTCGTTGATGACCTGCCAGCACCTTATGAGAAATCAAGTTTTGGGTTCGGGGGGAGTATGGTCGCAGGCTGAAACTTAAGGGAATTGACGGAAGGGCCCACCAGGCGGGGAACCGGCGGTTATTTGACTCCACCGGGAAAATTTCCGGCCCAAACCGGGGGGAAATAAAAAATTAAACTTTTTGTTTTTTGGGGGGGGGGGGGGGGCCCCTTTTTTTGGGGGGGGGCCCGCGG

> #101_18S

GGAAGCTGCTCAAGATTAAGCCATGCATGTCTAAGTATAAACTGCTTATACTGTGAAACTGCGAATGGCTCATTAAATCAGTTATAGTTTATTTGATGGTACCTTTACTCGGATAACCGTAGTAATTCTAGAGCTAATACGTGCGTAAATCCCGACTTCTGGAAGGGACGTATTTATTAGATAAAAGGCCAGCCGGGCTTGCCCGACTCTTGGCGAATCATGATAACTTCACGAATCGCACGGCTTCGTGCCGGCGATGTTTCATTCAAATTTCTGCCCTATCAACTTTCGATGGTAGGATAGAGGCCTACCATGGTGGTAACGGGTGACGGAGGATTAGGGTTCGATTCCGGAGAGGGAGCCTGAGAAACGGCTACCACATCCAAGGAAGGCAGCAGGCGCGCAAATTACCCAATCCCAACACGGGGAGGTAGTGACAATAAATAACAATACCGGGCATTTTTGTCTGGTAATTGGAATGAGTACAATCTAAATCCCTTAACGAGTATCCATTGGAGGGCAAGTCTGGTGCCAGCAGCCGCGGTAATTCCAGCTCCAATAGCGTATATTTAAGTTGTTGCAGTTAAAAAGCTCGTAGTTGGATTTCGGGTGGGTTGTAGCGGTCAGCCTTTGGTGAGTACTGCTACGGCCCACCTTTCTGCCGGGGACGTGCTCCTGGGCTTAACTGTCCGGGACACGGAATCGGCGAGGTTACTTTGAGTAAATTAGAGTGTTCAAAGCAAGCCTACGCTCTGAATACATTAGCATGGAATAACACGATAGGACTCTGGCTTATCTTGTTGGTCTGTAAGACCGGAGTAATGATTAAGAGGGACAGTCGGGGGCATTCGTATTTCATTGTCAGAGGTGAAATTCTTGGATTTATGAAAGACGAACTTCTGCGAAAGCATTTGCCAAGGATGTTTTCATTAATCAAGAACGAAAGTTGGGGGCTCGAAGACGATTAGATACCGTCGTAGTCTCAACCATAAACGATGCCGACTAGGGATTGGCAGGTGTTTCGTTGATGACCCTGCCAGCACCTTATGAGAAATCAAGCTTTTGGGTTCCGGGGGAAGTATGGTCGCAAGGCTGAAACTTAAAGGAATTGACGGAAGGCCCCACCAGGGGTGGAGCCTGCGGCTTATTTGACTCACACGGGAAAACTTACCAGGTCCAAACGGGGAAGAATGACAAATTGAAGCTCTTCCTGATCTGGGGGGAGGGGCATGCCGTCTTACCGTGGGGTGCCTGTCCAGGTGATTCCGGAACCAGCAACCCCAACCAAAAGAAACCCCTTCTCCCGGGGGCCGGGTCTTAAACCAATGGGGGTACCCATGAGGGGGGGGAAAACCCTCGGGAATCTTTTTTCCTGTCCCGCCCCCCCATTTTCTTGGGAACTCCCCGCGAGAGACGGGCTTTTTTATGTCTTGGGGCATATTCTTAT

> #127_18S

GGAAAGCTGTCTCAAGATTAAGCCATGCATGTCTAAGTATAAACTGCTTATACTGTGAAACTGCGAATGGCTCATTAAATCAGTTATAGTTTATTTGATGGTACCTTTACTCGGATAACCGTAGTAATTCTAGAGCTAATACGTGCGTAAATCCCGACTTCTGGAAGGGACGTATTTATTAGATAAAAGGCCAGCCGGGCTTGCCCGACTCTTGGCGAATCATGATAACTTCACGAATCGCACGGCTTCGTGCCGGCGATGTTTCATTCAAATTTCTGCCCTATCAACTTTCGATGGTAGGATAGAGGCCTACCATGGTGGTAACGGGTGACGGAGGATTAGGGTTCGATTCCGGAGAGGGAGCCTGAGAAACGGCTACCACATCCAAGGAAGGCAGCAGGCGCGCAAATTACCCAATCCCAACACGGGGAGGTAGTGACAATAAATAACAATACCGGGCATTTTTGTCTGGTAATTGGAATGAGTACAATCTAAATCCCTTAACGAGTATCCATTGGAGGGCAAGTCTGGTGCCAGCAGCCGCGGTAATTCCAGCTCCAATAGCGTATATTTAAGTTGTTGCAGTTAAAAAGCTCGTAGTTGGATTTCGGGTGGGTTGTAGCGGTCAGCCTTTGGTGAGTACTGCTACGGCCCACCTTTCTGCCGGGGACGTGCTCCTGGGCTTAACTGTCCGGGACACGGAATCGGCGAGGTTACTTTGAGTAAATTAGAGTGTTCAAAGCAAGCCTACGCTCTGAATACATTAGCATGGAATAACACGATAGGACTCTGGCTTATCTTGTTGGTCTGTAAGACCGGAGTAATGATTAAGAGGGACAGTCGGGGGCATTCGTATTTCATTGTCAGAGGTGAAATTCTTGGATTTATGAAAGACGAACTTCTGCGAAAGCATTTGCCAAGGATGTTTTCATTAATCAAGAACGAAAGTTGGGGGCTCGAAGACGATTAGATACCGTCGTAGTCTCAACCATAAACGATGCCGACTAGGGATTGGCAGGTGTTTCGTTGATGACCCTGCCAGCACCTTATGAGAAACCAAAGTTTTTGGCTTCCGGGGGGAGTATGGTCGCAGGCTGAAACTTAAAGGGATGGGACGGAAGGGCACACCAGGCGGGAGCCTGCGCTTATTGCTTCAACCGGGAAAACTATCAGCCCAGAACCGGGGAGCATGACGAATGAAGCTCTTTCTGGACCCGAGACGGCGCGCCAGACCACCAAACCGCTGCGTTAACGTTCCGGATCATGTTGCTACCCCCCTACTTCAATCTTTAGGAAACCCCCTCCCCGTGCGGGGAGATCCTAT

> *D.tertiolecta*_18S

GAGCCATGCATGTCTAGTATAAACTGCTTATACTGTGAAACTGCGAATGGCTCATAAATCAGTTATAGTTTATTTGATGGTACCTTTACTCGGATAACCGTAGTAATTCTAGAGCTAATACGTGCGTAAATCCCGACTTCTGGAAGGGACGTATTTATTAGATAAAAGGCCAGCCGGGCTTGCCCGACTCTTGGCGAATCATGATAACTTCACGAATCGCACGGCTTTATGCCGGCGATGTTTCATTCAAATTTCTGCCCTATCAACTTTCGATGGTAGGATAGAGGCCTACCATGGTGGTAACGGGTGACGGAGGATTAGGGTTCGATTCCGGAGAGGGAGCCTGAGAAACGGCTACCACATCCAAGGAAGGCAGCAGGCGCGCAAATTACCCAATCCCAACACGGGGAGGTAGTGACAATAAATAACAATACCGGGCATTTTTGTCTGGTAATTGGAATGAGTACAATCTAAATCCCTTAACGAGTATCCATTGGAGGGCAAGTCTGGTGCCAGCAGCCGCGGTAATTCCAGCTCCAATAGCGTATATTTAAGTTGTTGCAGTTAAAAAGCTCGTAGTTGGATTTCGGGTGGGTTGTAGCGGTCAGCCTTTGGTTAGTACTGCTACGGCCTACCTTTCTGCCGGGGACGAGCTCCTGGGCTTAACTGTCCGGGACTCGGAATCGGCGAGGTTACTTTGAGTAAATTAGAGTGTTCAAAGCAAGCCTACGCTCTGAATACATTAGCATGGAATAACACGATAGGACTCTGGCTTATCTTGTTGGTCTGTAAGACCGGAGTAATGATTAAGAGGGACAGTCGGGGGCATTCGTATTTCATGTCAGAGGTGAAATTCTTGGATTTATGAAAGACGAACTTCTGCGAAAGCATTTGCCAAGGATGTTTTCATTAATCAAGAAC

**SI 3** FASTA sequence of the ITS gene sequence of tested *Dunaliella* strains

>#006_ITS

AGCCCCGTAACTTTTGGATTCTTATACCGCTGCCCTTCAGAAACAACTGAGCGATTGCTGCCTAACCGTTTGGGTCCCTTGACGGGTAGCCGAGATAGCCCCTGCTCTTCAGCTGATCCTGATTCTTTGTTGGGGCCTGAATCACCCAAGCTCTGGAACAGCCAGGTCCACTTACGTTATTCCTCACATGAGGGAGTGGTTCCTCCTGTTAAAGAAGAAGGTTAAGGGTGGTTCCTTTGTTCAGCCAGACTGGATCTTCACCCAATAGTGAGGGAGGAGGCTATCATACTTTCTGGTTCGCATCTTTGTTTGTTGCGAAGTGCTACCCCGGTGAAGATTT

>#27_ITS

TGATGATTCACGGAATTCTGCAATTCACACTACGTATCGCATTTCGC

>#37_ITS

TCTTAGGTTTTGGAGGGCCAAGCCCATGGTCCCAAGCCAACAACTAGAAATAGCGAGCGATTGCTGCCTACCCAGTTGCGGCCCTTGACGGGTCCTTTGAGCTAGCCTCTGCTCTTCAGCTGATCCAGAGCCTTTGTTGGGGCAGTGAAGCACCCAAGCTCTGGAACAGCCAGGTCCACTAACATTACTCCTCACATGGGGGAGTGGTTAGTGAGATTAACCCGACGCTGAGGCAAACATGCCCTTGGCCGAAGCCGCGGACGCAATTTGCGTTCAAAGATTTGATGATTCACGGAATTCTGCAATTCACACTACGTATCGCATTTCGCTGCGCCTCTTCACCGAAGCGCCA

>#83_ITS

GCTTCGGTCGAGAAAAGAAGGGTTCCTGTTTGAGGGCCAAGCCCATGGTCCCAAGCCAACAACTAGAAATAGCGAGCGATTGCTGCCTACCCAGTTGCGGCCCTTGACGGGTCCTTGAGCTAGCCTCTGCTCTTCAGCTGATCCAGAGCCTTTGTTGGGGCAGTGAAGCACCCAAGCTCTGGAACAGCCAGGTCCACTAACATTACTCCTCACATGGGGGAGTGGTTAGTGAGATTAACCCGACGCTGAGGCAAACATGCCCTTGGCCGAAGCCGCGGACGCAATTTGCGTTCAAAGATTTGATGATTCACGGAATTCTGCAATTCACACTACGTATCGCATTTCGCTGCGTTCTTAATCA

>#96_ITS

GAGCTCAGGTCGAGAAAAAAAGAGGTTCCTGTTTGAGGGCCGAGCCCATGGTCCCAAGCCAACAACTAGAAATAGCAAGCGTGAGCTGCCTACCCAGTTGCGGCCCTTGACGGGTCCTTGAGCTAGCCTCTACTCTTCAGCTGATCCAGAGCTTTTGCTAAGCGCAATGAAGCACTCAAGCTCTGGAACAGCCAGGTCCACTTACCCGCTTCCACTATGGGAGGAGGGGAGTGAGATTAACCCGACGCTGAGGCAAACATGCCCTCAGCCGAAGCCTTGGGCGCAATTTGCGTTCAAAGATTTGATGATTCACGGAATTCTGCAATTCACACTACGTATCGCATTTCGCTGCGTTTTA

>#*D.tertiolecta*_ITS

TCTGCTTGAGCTCAGGTCGAGAAAAAAAGAGGTTCCTGTTTGAGGGCCGAGCTCATGGTCCCAAGCCAACAACTAGAAATAGCAAGCGTGAGCTGCCTACCCAGTTGCGGCCCTTGACGGGTCCTTGAGCTAGCCTCTACTCTTCAGCTGATCCAGAGCTATTGCCAAGCGCAATGAAGCACTCAAGCTCTGGAACAGCCAGGTCCACTTACCCGCTTCCACTATGGGAGGAGGGGAGTGAGATTAACCCGACGCTGAGGCAAACATGCCCTCAGCCGAAGCCTTGGGCGCAATTTGCGTTCAAAGATTTGATGATTCACGGAATTCTGCAATTCACACTACGTATCGCATTTCGCTGCGTTCTTAAT

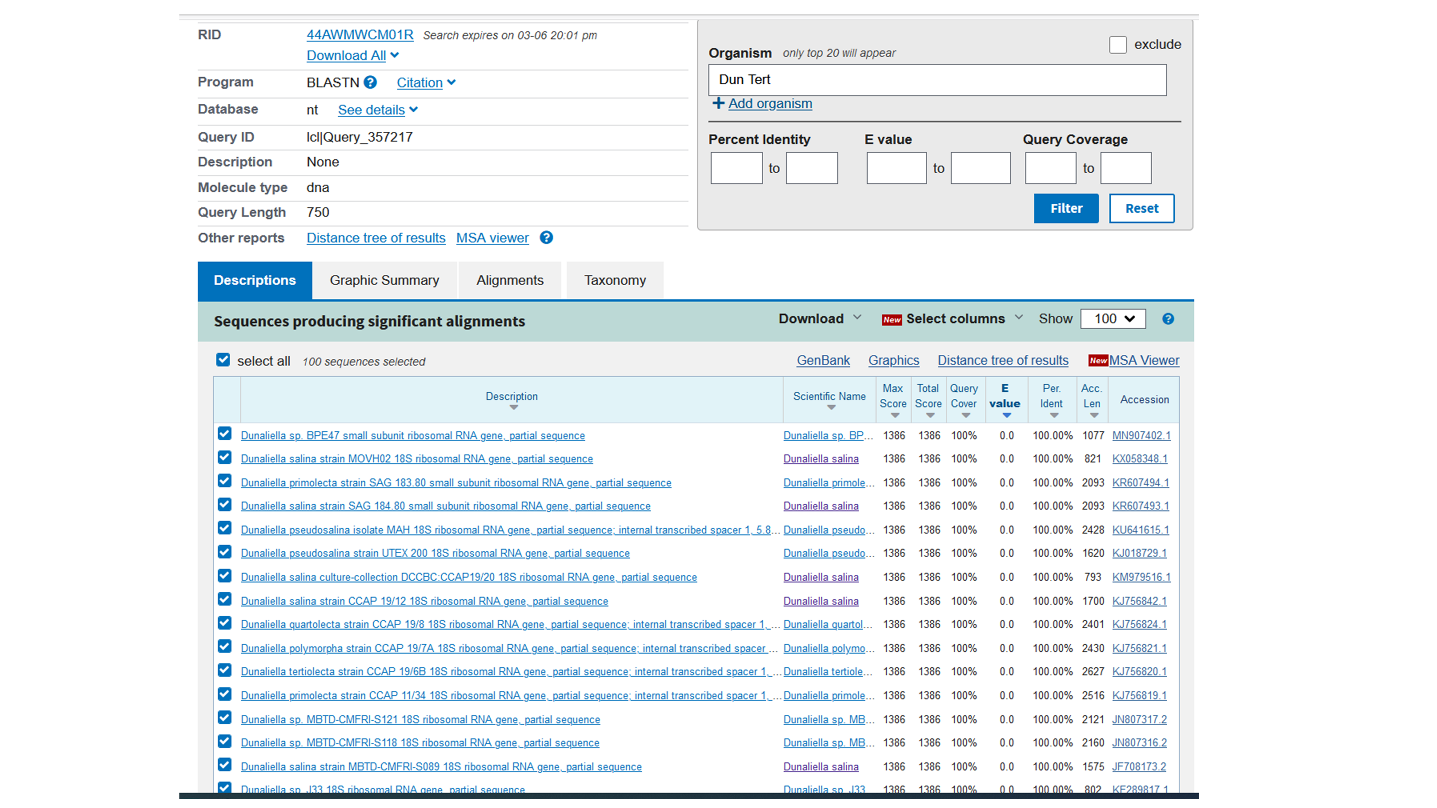

**SI 4** 18S gene sequence of *D. tertiolecta* was searched against the GenBank database by the BLASTn algorithm resulting in many hits with a query cover and percentage identity of 100%. All hits identify this strain as a *Dunaliella* strain, but did not allow identification of the exact species.

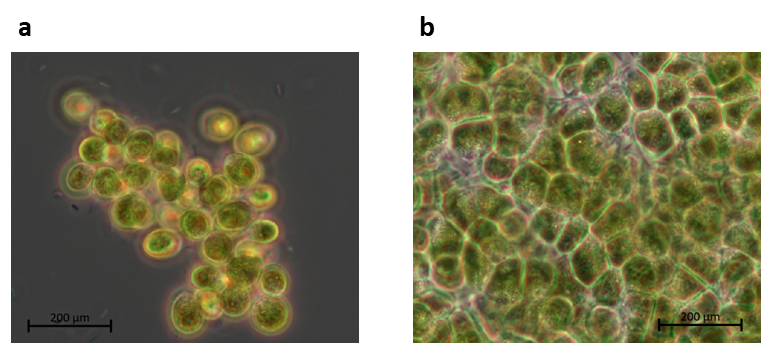

**SI 5** Bright field images (100x) of isolate #96 when cultivated in a) 1.5 M and b) 2 M NaCl containing modified Johnson medium (pH=7.5). The higher the salt concentration, the earlier the cells start to accumulate and the larger the formed palmella structures.
